# QuImputer: an LD-informed QUBO/Ising formulation of genotype imputation with quantum-hardware execution

**DOI:** 10.64898/2026.09.26.754587

**Authors:** Minwoo Kim, Tae-Ho Lee

## Abstract

Genotype imputation offers a biological application for optimization methods that may eventually run on larger quantum computers. We developed QuImputer to encode allele-frequency priors and signed linkage disequilibrium (LD) in a reduced Ising/QUBO objective, with only missing genotypes represented as optimization variables. The same objective can be passed to exhaustive classical search, quantum approximate optimization algorithm simulation, or quantum hardware. We evaluated the formulation on 100 rice accessions across 12 chromosomes, estimating LD and allele frequency from the same post-masking observed matrix supplied to Beagle 5.4. Evaluation was restricted to known truth at sites with exact dosage coding and an alternate allele that was not the major allele, with MAF below 0.50, yielding 1.88 billion scored genotypes. Beagle achieved higher rare-ALT sensitivity (2.52% versus 0.54%), F1 (0.047 versus 0.011), Matthews correlation coefficient (0.088 versus 0.044), and pooled hard-call dosage R^2^ (0.231 versus 0.014). QuImputer produced fewer false ALT calls, with Type I error of 0.038% versus 0.223%. Analyses of a chromosome-1 subset showed limited local LD support for many masked rare-ALT carriers. In five small IBM Fez validation problems, the selected minimum-energy outputs matched unique all-reference exact optima; the pooled frequency of those optima was 1.21-2.63%. Further IBM Fez executions covered eleven sparse instances with 12-97 variables and a separate 120-variable instance, retaining all measurements. Selected outputs recovered the exact optima in eight instances through 49 variables. Larger instances remained below the all-reference baseline; noiseless comparisons distinguished finite-shot sampling from deviations in hardware output. QuImputer provides an LD-informed formulation and a route to quantum-hardware execution, together with an empirical assessment of the limitations of the evaluated configuration. These results establish a basis for studying richer information models and alternative classical or quantum solvers.

## 1. Introduction

Accurate genotype data are central to association genetics and crop improvement, but the cost of deep sequencing limits the number of individuals that can be characterized. Agricultural breeding programs therefore frequently use low-coverage genotyping-by-sequencing (GBS), which can produce extensive and spatially clustered missingness (Elshire et al., 2011; Swarts et al., 2014; Torkamaneh et al., 2016). Imputation is used to infer these unobserved genotypes from population haplotypes, allele frequencies, and linkage disequilibrium (LD).

Hidden Markov model (HMM) methods derived from the Li-Stephens haplotype-copying framework are established approaches for genotype imputation (Li and Stephens, 2003). Beagle, IMPUTE2, Minimac, STITCH, and GLIMPSE exploit chromosome-scale haplotype structure or large reference panels and generally perform well when informative markers and compatible haplotypes are available (Browning and Browning, 2007; Howie et al., 2009; Davies et al., 2016; Das et al., 2016; Browning et al., 2018; Rubinacci et al., 2021). Other approaches, such as LinkImpute, use local LD and nearest-neighbour information for non-model organisms (Money et al., 2015). Under severe and spatially clustered missingness, however, evidence supporting a rare allele may be limited, and probabilistic calls may be influenced strongly by the more frequent population state. The extent of this behaviour depends on marker density, reference composition, population structure, and software configuration and should be measured directly in each benchmark rather than assumed.

A complementary formulation is to treat a window of missing genotypes as a combinatorial optimization problem. If allele-frequency priors and pairwise LD relationships define an energy landscape, genotype assignment can be expressed as an Ising Hamiltonian or quadratic unconstrained binary optimization (QUBO) problem (Lucas, 2014; Glover et al., 2019). This representation is solver-agnostic: the same QUBO can be evaluated by exhaustive classical enumeration, classical heuristics, a quantum simulator, or a quantum processing unit (QPU). For gate-based optimization, the quantum approximate optimization algorithm (QAOA; Farhi et al., 2014) is a variational quantum algorithm (Cerezo et al., 2021) that has been studied extensively for combinatorial optimization (Blekos et al., 2024; Abbas et al., 2024). Quantum optimization has previously been explored for biological problems including protein folding, de novo sequence assembly, and epistasis-oriented SNP selection (Perdomo-Ortiz et al., 2012; Sarkar et al., 2021; Boev et al., 2021; Hoffmann et al., 2024). Date et al. (2019) listed genotype imputation as a candidate QUBO application, but, to our knowledge, a complete genotype-imputation implementation with gate-based QPU validation has not previously been reported.

Here we present QuImputer, a reduced LD-informed QUBO/Ising formulation in which only missing SNPs become optimization variables. Our aim is to make the imputation objective explicit and portable across classical and quantum optimization methods. We characterize its binary, dosage and rare-ALT error profiles against Beagle using information available after masking, measure the effect of window size on performance and exhaustive classical cost, and test selected circuits on a superconducting QPU. Diagnostic and secondary analyses examine local information, parameter sensitivity and alternative information conditions. This study establishes the formulation and its execution path while documenting the capabilities and limitations of the current implementation.

## 2. Methods

### 2.1 Dataset and masking design

Rice is a well-characterized model for crop diversity and association genetics (McNally et al., 2009; Huang et al., 2010). From the 3,000 Rice Genomes Project dataset (3,024 accessions and approximately 28.8 million SNPs), we ranked all accessions by per-sample genotype missingness (VCFtools --missing-indv, evaluated over 28,761,754 sites; Danecek et al., 2011) and retained the 100 accessions with the lowest missingness as the truth panel. No additional filter was applied, and no tie occurred at the selection boundary (100th F_MISS = 0.0634, 101st = 0.0640). The retained panel had per-sample missingness of 0.021-0.063 and mean sequencing depth of 7.1-28.5x; depth is reported as a descriptive statistic of the panel and was not used as a selection criterion (The 3,000 Rice Genomes Project, 2014; Wang et al., 2018). The identifiers, per-sample quality-control metrics and variety-group assignments of these 100 accessions are provided as Supplementary Data 1. Selecting on missingness produced a panel that is strongly skewed towards lineages close to the reference assembly: 84.0% of the panel belongs to the japonica groups against 27.97% of the 3,000-accession collection, whereas indica accessions, which make up 58.10% of the collection, contribute 7.0%, and the aromatic groups are absent entirely (Supplementary Table S14).

To emulate a deliberately severe low-coverage GBS setting, we masked 60% of REF genotypes and 90% of ALT-containing genotypes using random seed 42 and initiated 50-200-SNP blackout blocks with probability 0.5%. The 60/90% rates were heuristic stress-test settings rather than estimates fitted to this dataset. Blackout blocks were included to represent extended regions with little observed information. A secondary allele-frequency-independent 70% masking analysis is reported in Supplementary Table S2. Because that analysis uses the complete-panel information condition defined in Section 2.3, it evaluates sensitivity to masking structure but does not estimate the primary performance margin.

Beagle 5.4 (Browning et al., 2021) was run on the same masked matrix with default parameters and without an external genetic map. This represents a portable baseline rather than a rice-optimized upper bound; evaluation with tuned Beagle settings and empirical genetic maps remains future work.

### 2.2 QUBO/Ising formulation

Each SNP state was encoded as an Ising spin s_i_ in {−1,+1}, where −1 denotes homozygous reference and +1 denotes an ALT-carrier state. Heterozygous and homozygous ALT genotypes were therefore combined for binary concordance and classification metrics. This binary representation is suited to testing the formulation in predominantly homozygous rice, but it does not distinguish heterozygous dosage. Supplementary Table S1 documents the original chromosome-1 encoding summary; those counts use the allele-index-sum coding that the primary evaluation later replaced and are retained as a description of the encoding rather than as evaluation quantities.

The energy function was

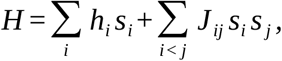

where the local bias for a missing SNP *i* was

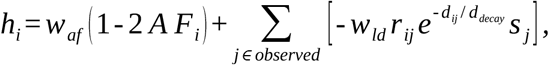

and pairwise couplings between missing SNPs were

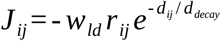

Here *A F*_*i*_ is allele frequency, *r*_*ij*_ is signed Pearson correlation, *d*_*ij*_ is physical distance, and *d*_*decay*_ = 10, 000 bp. Only missing SNPs became QUBO variables; observed SNPs contributed through bias terms. Couplings with *r*^2^ < 0.01 were discarded in the classical benchmark. Series A hardware instances used *r*^2^ thresholds of 0.01 for 12 and 15 qubits, 0.1 for 18 qubits, and 0.3 for 25-97 qubits (Supplementary Table S10). The Ising variables were converted to binary QUBO variables by *s*_*i*_ = 2 *x*_*i*_ - 1.

The separate 120-variable hardware instance used an LD r^2^ threshold of 0.5 and a 200-SNP window (Supplementary Table S16).

For an upper-triangular QUBO matrix, the equivalent binary objective and coefficients are

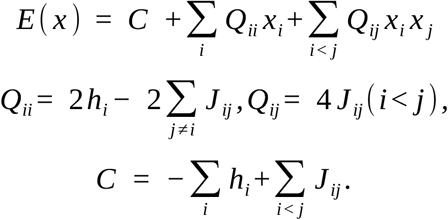

Here J is extended symmetrically in the diagonal formula, and the constant C does not affect the minimizer. The archived hardware Hamiltonians use Pauli-Z values z_i_=1−2x_i_=−s_i_; their local fields therefore have the opposite sign, while pairwise coefficients are unchanged.

Chromosomes were divided into non-overlapping windows of *w* SNPs. Each window was solved independently, so LD across window boundaries was not directly represented. QuImputer returns REF or ALT-carrier assignments, not calibrated genotype probabilities or heterozygous dosage.

### 2.3 Primary and secondary information conditions

For the **primary observed-data condition**, *A F*_*i*_ and *r*_*ij*_ were estimated from the post-masking observed genotype matrix. Thus, masked genotypes used for scoring were not used directly to construct evaluation-time QuImputer statistics. This is the same genotype matrix supplied to Beagle, although the two algorithms use that information differently.

A secondary complete-panel condition estimated LD and allele frequency from the complete pre-masking genotypes of the same 100 samples. It includes information about genotypes hidden for scoring and serves as a developmental diagnostic. Secondary analyses are labelled throughout. Their archived evaluation definition differs from the corrected primary definition, so whole-genome primary and secondary values in Supplementary Table S8 are presented as separate analysis records rather than a matched estimate of the information-source effect. A five-seed masking analysis on 100,000 chromosome-1 SNPs under this secondary condition is reported in Supplementary Table S3.

A 30-combination parameter grid was evaluated on a 1,000,000-SNP development analysis under the complete-panel condition. The selected weights, w_ld = 0.5 and w_af = 0.3, were applied unchanged to the primary observed-data evaluation. They are development choices whose generalizability requires an independent observed-data assessment (Supplementary Table S4; Supplementary Figure S3). A common positive rescaling of both weights leaves the exact argmin unchanged; the grid therefore contains 21 distinct weight ratios. The Table 5 validation problems and all October hardware instances were constructed under the complete-panel condition, with LD and allele frequency computed from pre-masking genotypes and the masked matrix supplying observed states and the missing-variable set. They are separate from the primary observed-data benchmark.

### 2.4 Solver hierarchy and quantum hardware

For a fixed instance, each solver acts on the same reduced QUBO objective or its equivalent Ising representation (Figure 1). The whole-genome evaluation and selected hardware experiments use the information conditions specified in Section 2.3.

**Figure 1.**
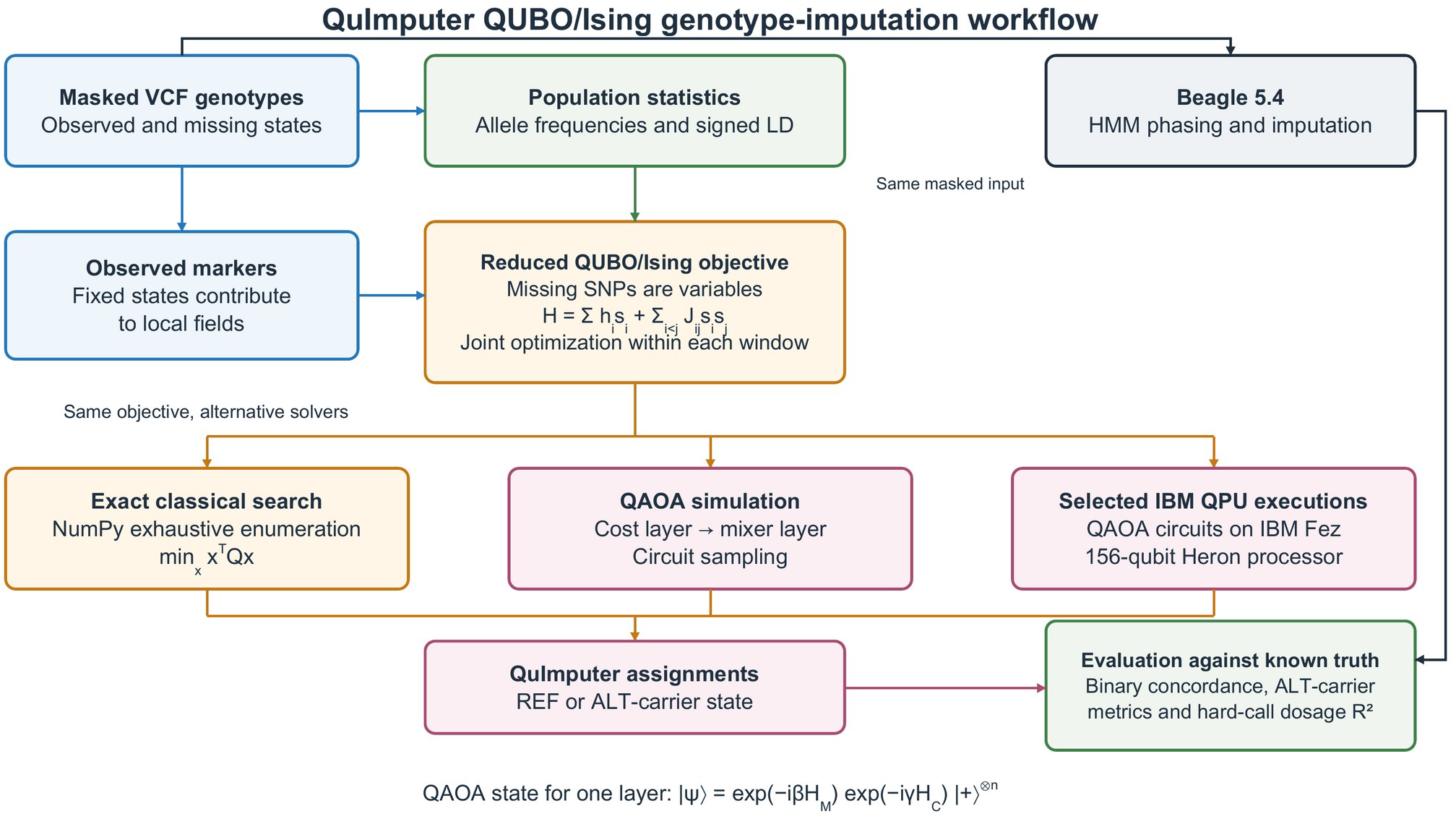
QuImputer QUBO/Ising genotype-imputation workflow and solver hierarchy. VCF genotypes are summarized through a signed LD matrix; observed markers contribute bias terms and missing SNPs become variables in a reduced Ising/QUBO problem. The same QUBO can be solved by exhaustive NumPy search, QAOA simulation, or selected IBM QPU executions. Beagle 5.4 provides an HMM-based phasing and imputation comparator using the same masked input matrix in the primary benchmark. Both outputs are scored against held-out genotypes. Primary LD and allele-frequency estimates are obtained from the post-masking observed matrix; the complete-panel condition is used in designated secondary and hardware analyses. The QAOA state applies the cost layer before the mixer layer, and optimization is joint within each window.

1. **Exhaustive classical solver**. A vectorized NumPy implementation enumerated all 2^*n*^ states in chunks and returned the guaranteed global minimum. This solver generated the full *w* = 25 and *w* = 30 benchmark results.
2. **QAOA simulator**. The five validation problems in Table 5 were executed on 18 September 2026 using one-layer QAOA with depth *p* = 1 and Qiskit Aer SamplerV2. Each problem used 10 equally spaced gamma values from -π/2 to π/2 and 10 beta values from -π/4 to π/4, including both endpoints, with gamma as the outer loop. Each of the 100 circuits was sampled 4,096 times, giving 409,600 shots per problem per backend. The problems were selected from a 20-SNP window at the start of chromosome 1 using complete-panel LD and allele frequencies, with the LD threshold set to 0.01.
3. **IBM QPU**. The same logical circuits were executed on IBM Fez, a 156-qubit Heron processor, with 4,096 shots per circuit. Both backends used transpiler optimization level 1. No explicit sampler or transpiler seed was set. Qiskit 2.3.1, Aer 0.17.2 and IBM Runtime 0.46.1 were used; instance identities, software versions, job identifiers and selected outputs are provided in Supplementary Table S15 and the supplementary data.

For both backends, the final output was the lowest-energy string observed across all 100 circuits. Energy ties retained the first string in the recorded counts iteration order within a circuit and the first candidate in grid order across circuits, using strict less-than comparisons. Measured strings were reversed to logical variable order before evaluation, with 0 denoting REF and 1 denoting ALT carriage. Exact solutions and reference genotypes were used for evaluation and recording, not to tune the parameter grid, select the output, or correct it. Concordance is the number of matching variables divided by the number of active variables. Pooled exact-optimum frequency is the number of shots equal to the exact bitstring divided by 409,600; it combines different parameter settings and is distinct from selected-output concordance.

On 6 October 2026, we executed eleven instances matching the archived G3 coefficients and binary truth (series A, 12-97 variables), together with one newly documented 120-variable instance (B120), on ibm_fez. The retained selection procedure used the first 100,000 chromosome-1 SNPs for A and the first 50,000 SNPs with 200-SNP windows for B120; genomic identities and parameter settings are provided in Supplementary Tables S13 and S16. These are new dated executions, separate from the September Table 5 experiment. A coefficients match the archived objectives, whereas B120 was selected using the retained historical procedure without coefficient-level confirmation against the March execution.

Each October instance used one-layer QAOA: Hadamard initialization, RZ(2γh_i_), CX-RZ(2γJ_ij_)-CX, and RX(2β), in the circuit convention z=1-2x. Both parameter grids included their endpoints: γ in [-π/2, π/2] and β in [-π/4, π/4], with gamma as the outer loop. A used 5×5 settings and B120 used 10×10 settings; each circuit had 4,096 shots. Circuits were transpiled at optimization level 0 with seed 42. Gate and measurement twirling and dynamical decoupling were disabled. Qiskit 2.3.1 and IBM Runtime 0.46.1 were used, with one Sampler job per instance. Core instance and execution records are provided in the supplementary data; full records are available on request. IBM-reported QPU usage totalled 431 s across 375 circuits and 1,536,000 shots on an Open-plan account.

October outputs used the same minimum-observed-energy selection principle as Table 5, with energies accumulated in logical-variable order, each local-field term followed by its pairwise terms. Strict comparisons retained the first observed string on ties and the first circuit in grid order. Truth and exact references were used only after output selection. Concordance with biological truth excludes variables with missing original truth; concordance with the exact solution includes all variables. Exact optima and noiseless circuit probabilities were computed by enumerating connected components of the coupling graph. Expected optimum counts sum 4,096p_c_ across circuits, where p_c_ is each circuit’s exact optimum probability. Under independent ideal shots, the probability of observing no optimum is the product of (1-p_c_)^4,096^. A separate descriptive comparison used 500 noiseless finite-shot replicates per instance, seed 42, with the identical energy accumulation and selection rule (Supplementary Table S16).

The *w* = 20 timing benchmark used a 64-core, 2-TB server. Whole-genome *w* = 25 runs used 100 Cray XC40 nodes per chromosome (2,400 MPI ranks), and *w* = 30 used 200 nodes per chromosome (4,800 MPI ranks). A separate identical-hardware 100,000-SNP benchmark measured *w* = 20, *w* = 25, and *w* = 30 on the 64-core server.

### 2.5 Evaluation metrics and analysis universes

The primary evaluation includes artificially masked genotypes with non-missing original truth, excludes sites with corrected alternate-allele frequency above 0.50, and retains sites whose original dosage coding exactly represents non-reference copy number. The MAF intervals are [0,0.01), [0.01,0.05), [0.05,0.10) and [0.10,0.50); positions at exactly MAF = 0.50 are excluded from the overall total. These rules give 1,877,517,664 scored genotypes, including 323,536,811 in the MAF 1–10% rare-ALT analysis. Corrected truth-derived allele frequency defines evaluation bins and exclusions only; the primary predictor uses post-masking observed statistics. Supplementary Table S7c documents staged changes from the previous evaluation definition.

The original implementation obtained dosage by summing VCF allele indices. We audited this coding against non-reference copy number, defined as the number of alleles whose index is greater than zero. The exact-coding set includes sites with a single non-deletion ALT entry and sites at which the only observed nonzero allele index is 1. Sites with other observed nonzero indices are excluded from primary scoring. In a diagnostic of 4,000 windows from the first 100,000 SNPs on chromosome 1, correcting the coding changed threshold-based edge inclusion for 2 of 1,200,000 marker pairs; 99.95% of windows retained the same edge set. This statistic describes edge inclusion, not equality of all retained coupling coefficients. The reported primary results re-score the existing predictions at exactly coded sites; they do not represent a new imputation run with every predictor statistic recomputed under corrected coding.

Binary concordance compares REF versus ALT-carrier states at the primary evaluation positions, with dosage 0 mapped to REF and dosage 1 or 2 mapped to ALT-carrier. Pooled hard-call dosage R^2^ is the squared Pearson correlation between truth dosage and predicted dosage over these same 1,877,517,664 positions, reconstructed from per-chromosome sufficient statistics (Supplementary Table S9). QuImputer predictions are encoded as 0 or 2. Beagle dosage is derived from its hard GT calls: the reference-free runs used here did not populate DS, and reruns with gp=true, or gp=true together with ap=true, left GT calls unchanged. The statistic is pooled across positions and samples, rather than averaged across per-SNP correlations. Applying the 0/2 mapping to Beagle and to truth provides a matched binary comparison and an oracle recoding reference, respectively.

Rare-ALT classification uses the MAF 1–10% primary evaluation set. Sensitivity is TP/(TP+FN), Type I error is FP/(FP+TN), precision is TP/(TP+FP), F1 is 2TP/(2TP+FP+FN), and balanced accuracy is the mean of sensitivity and specificity. Matthews correlation coefficient (MCC) summarizes the four confusion counts. Type II error is 1−sensitivity.

Gene-level analysis used RAP-DB IRGSP-1.0 representative gene features (Sakai et al., 2013). Genes overlapping data SNPs with at least three rare SNPs and at least 50 masked rare-genotype evaluations were retained. The 29,437 eligible genes contribute 58,390,209 evaluations to the gene-region aggregate in Table 2; overlapping genes were counted independently. Per-gene Type II comparisons additionally required at least 10 true ALT-carrier evaluations, yielding 28,187 genes. Differences within ±0.1 percentage points were classified as tied or similar; counts labelled as lower Type II error exceed this tolerance (Supplementary Figure S2).

## 3. Results

### 3.1 Primary whole-genome performance

The primary evaluation comprises 1,877,517,664 scored genotypes, including 323,536,811 in the MAF 1–10% analysis (Methods). At w=25, QuImputer had slightly higher concordance than Beagle at MAF <1% and 1–5%, whereas Beagle had higher concordance at MAF 5–10% and 10–50% (Table 1; Figure 2). Both methods fell below the all-REF baseline at MAF <1% and 1–5%, and both exceeded it in the two higher-frequency bins. Across all scored positions, the excess over all-REF was 0.0218 percentage points for QuImputer and 0.2555 for Beagle. Because reference calls dominate the low-MAF bins, direct ALT-carrier metrics are needed to assess recovery of rare variants (Section 3.2). Per-chromosome values are given in Supplementary Table S6 and Supplementary Figure S1.

**Table 1.** Whole-genome performance under the primary observed-data condition.

| Metric / MAF bin | QuImputer $w=25$ | QuImputer $w=30$ | Beagle 5.4 | Evaluated N |
| --- | --- | --- | --- | --- |
| Binary concordance, MAF $<1\%$ | 0.999012 | 0.999010 | 0.998992 | 1,508,245,134 |
| Binary concordance, MAF 1-5% | 0.956730 | 0.956665 | 0.955750 | 276,033,817 |
| Binary concordance, MAF 5-10% | 0.884728 | 0.884655 | 0.885791 | 47,502,994 |
| Binary concordance, MAF 10-50% | 0.697346 | 0.698522 | 0.798750 | 45,735,719 |
| Overall binary concordance | 0.982555 | 0.982571 | 0.984892 | 1,877,517,664 |
| Excess over all-REF baseline, MAF $<1\%$ | -0.0007 pp | -0.0008 pp | -0.0026 pp | 1,508,245,134 |
| Excess over all-REF baseline, MAF 1-5% | -0.0131 pp | -0.0196 pp | -0.1111 pp | 276,033,817 |
| Excess over all-REF baseline, MAF 5-10% | +0.0268 pp | +0.0195 pp | +0.1331 pp | 47,502,994 |
| Excess over all-REF baseline, MAF 10-50% | +0.9677 pp | +1.0854 pp | +11.1081 pp | 45,735,719 |
| Excess over all-REF baseline, overall | +0.0218 pp | +0.0234 pp | +0.2555 pp | 1,877,517,664 |
| Dosage R2, MAF $<1\%$ | 0.000000 | 0.000000 | 0.000000 | 1,508,245,134 |

| Metric / MAF bin | QuImputer w=25 | QuImputer w=30 | Beagle 5.4 | Evaluated N |
| --- | --- | --- | --- | --- |
| Dosage R2, MAF 1-5% | 0.000775 | 0.000788 | 0.001779 | 276,033,817 |
| Dosage R2, MAF 5-10% | 0.003268 | 0.003515 | 0.033058 | 47,502,994 |
| Dosage R2, MAF 10-50% | 0.013642 | 0.015483 | 0.353051 | 45,735,719 |
| Pooled dosage R2 | 0.014339 | 0.016068 | 0.231099 | 1,877,517,664 |

The all-REF baseline assigns REF at every evaluated position. Its concordance is 0.999018, 0.956861, 0.884460 and 0.687669 in the four MAF bins, and 0.982337 overall. Applying the same 0/2 recoding to Beagle gives pooled dosage $R^2 = 0.211538$ ; applying it to truth gives the oracle recoding reference $R^2 = 0.888768$ . All entries use the primary evaluation definition in Methods.

**Figure 2.**
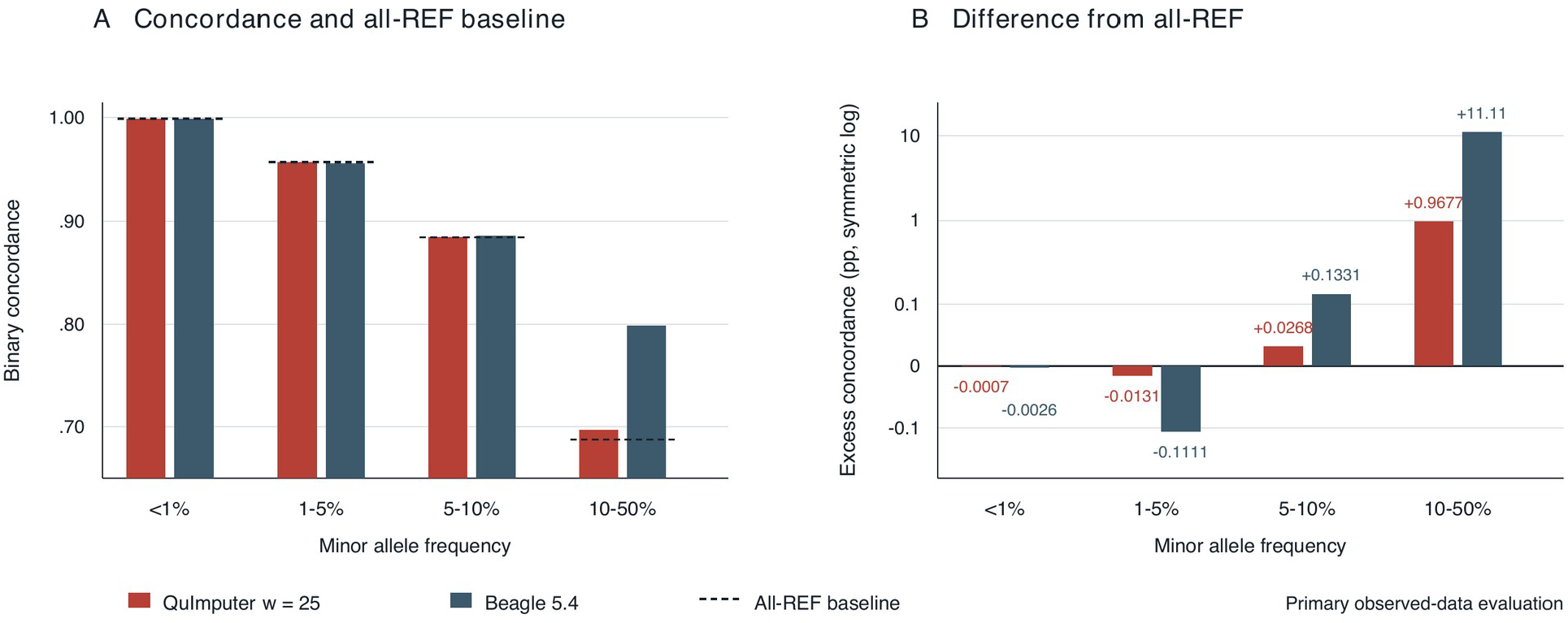
MAF-binned concordance in the primary observed-data evaluation. (A) QuImputer (w=25), Beagle and the all-REF baseline. (B) Excess over all-REF on a symmetric logarithmic scale. Both methods are below this baseline at MAF <1% and 1–5% and above it at MAF 5–10% and 10–50%. Table 1 gives exact values and counts; Figure 3 shows direct ALT-carrier metrics.

Pooled hard-call dosage R^2^ was 0.014339 for QuImputer at w=25 and 0.016068 at w=30, compared with 0.231099 for Beagle. Binary representation alone does not account for this gap: recoding Beagle to 0/2 gives R^2^ = 0.211538, while recoding truth to 0/2 gives an oracle reference of 0.888768. The largest bin-level difference occurred among common variants, with R^2^ = 0.013642 for QuImputer and 0.353051 for Beagle. Values at MAF <1% rounded to 0.000000 for both methods; at MAF 1–5% they were 0.000775 and 0.001779. These are pooled hard-call statistics and do not assess calibrated genotype probabilities.

### 3.2 Rare-ALT classification and gene-level heterogeneity

Rare-ALT classification showed different calling profiles (Table 2; Figure 3). Genome-wide, Beagle detected 4.7 times more true carriers, with sensitivity of 2.52% compared with 0.54% for QuImputer, while issuing 1,120,354 rather than 210,470 ALT calls. QuImputer made 93,515 true-positive and 116,955 false-positive calls; Beagle made 438,435 and 681,919. QuImputer therefore had lower Type I error (0.038% versus 0.223%) and higher genome-wide precision (44.43% versus 39.13%). Beagle had higher F1 (0.0474 versus 0.0106) and MCC (0.0882 versus 0.0442). In gene regions, Beagle also had higher precision (45.36% versus 44.29%), while QuImputer retained lower Type I error. An alternative evaluation that recoded ALT-major sites as minor-allele carriers gave the same qualitative sensitivity and precision ordering (Supplementary Table S7c). Each method is represented by one operating point; the comparison does not establish a precision–recall frontier.

**Table 2.** Rare ALT-carrier classification under the primary observed-data condition (w=25). Raw genome-wide and gene-region confusion counts are provided in Supplementary Table S7a.

| Analysis level / metric | QuImputer | Beagle 5.4 | Interpretation |
| --- | --- | --- | --- |
| Genome-wide, MAF 1-10% (N=323,536,811) |  |  |  |
| Sensitivity (1 - Type II error) | 0.54% | 2.52% | Beagle higher |
| Type I error | 0.038% | 0.223% | QuImputer lower |
| Precision | 44.43% | 39.13% | QuImputer higher |
| F1 score | 0.0106 | 0.0474 | Beagle higher |
| Matthews correlation coefficient | 0.0442 | 0.0882 | Beagle higher |
| Balanced accuracy | 0.5025 | 0.5115 | Beagle higher |
| Annotated gene regions (58,390,209 calls; 29,437 filtered genes) |  |  |  |
| Sensitivity (1 - Type II error) | 0.75% | 2.96% | Beagle higher |
| Type I error | 0.056% | 0.212% | QuImputer lower |
| Precision | 44.29% | 45.36% | Beagle higher |
| F1 score | 0.0148 | 0.0556 | Beagle higher |
| Matthews correlation coefficient | 0.0519 | 0.1047 | Beagle higher |
| Balanced accuracy | 0.5035 | 0.5137 | Beagle higher |
| Genes with Type II error lower by >0.1 pp (28,187 genes) | 2,395 (8.50%) | 8,503 (30.17%) | 17,289 tied or within 0.1 pp (61.34%) |

**Figure 3.**
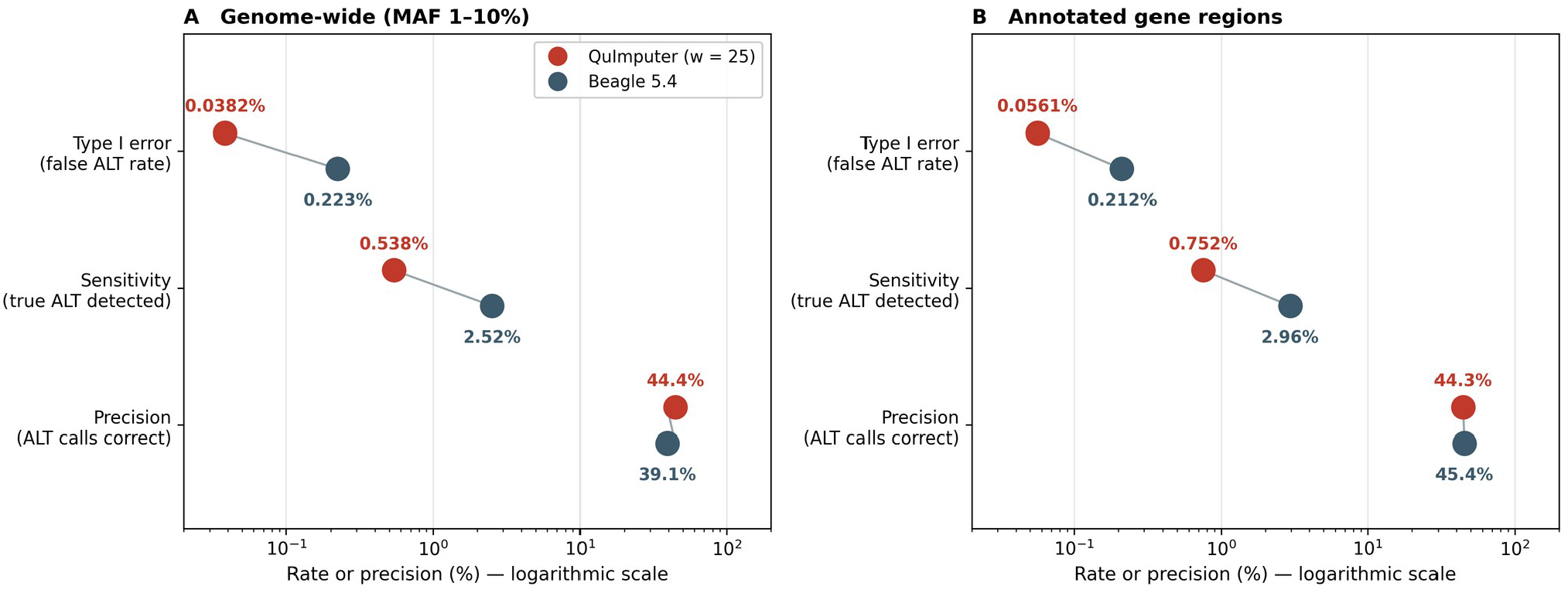
Rare-ALT error profiles in the primary observed-data evaluation. Type I error, sensitivity and precision are shown on a logarithmic axis for (A) genome-wide MAF 1–10% positions (N=323,536,811) and (B) annotated gene regions (N=58,390,209). QuImputer produces fewer false ALT calls, while Beagle recovers more true carriers. The genome-wide precision margin for QuImputer is not retained in gene regions. Raw confusion counts are given in Supplementary Table S7a.

Among the 28,187 genes eligible for per-gene comparison, QuImputer had lower Type II error by more than 0.1 percentage points in 2,395 (8.50%), Beagle in 8,503 (30.17%), and 17,289 (61.34%) were tied or within the 0.1-point tolerance. The w=30 analysis showed the same qualitative pattern (Supplementary Table S7b; Supplementary Figure S2).

Individual genes at the favourable tail of this distribution illustrate the heterogeneity. Applying a filter that required no contribution from sites at which the alternate allele is the major allele, at least 50 true carrier evaluations, lower Type II error for QuImputer, and no increase in false positives, 1,802 genes qualified. Among the highest ranked, Os05g0115700 (similar to UDP-galactose transporter related protein) contained 483 true carriers of which QuImputer recovered 36 with 1 false positive, against 0 recovered and 3 false positives for Beagle; Os10g0513900 (cytochrome P450 family protein) contained 144 true carriers with 18 recovered and no false positives, against 2 recovered and 10 false positives; and Os07g0166700 (LRR receptor-like kinase family protein) contained 50 true carriers with 7 recovered and no false positives, against 0 recovered and 1 false positive. These cases lie at the extreme of a distribution in which Beagle has the lower Type II error in 30.17% of genes against 8.50% for QuImputer. They are exploratory and post hoc, the annotations are those of RAP-DB rather than evidence of biological relevance, and the pattern of simultaneously higher recall and fewer false positives seen here is not a general property of the QuImputer-favoured tail.

### 3.3 Effect of optimization-window size

Increasing the window from 25 to 30 SNPs produced small sensitivity gains with lower precision. Binary concordance changed by −0.0065 and −0.0073 percentage points at MAF 1–5% and 5–10%, by +0.1176 among common variants, and by +0.0016 overall; pooled dosage R^2^ increased by 0.001728 (Table 3). Genome-wide rare-ALT sensitivity rose from 0.54% to 0.62%, Type I error from 0.038% to 0.050%, and precision fell from 44.43% to 41.35%. MCC increased from 0.0442 to 0.0452 and F1 from 0.0106 to 0.0121. Gene-region metrics showed the same trade-off. These two window sizes provide a local sensitivity comparison, rather than an estimate of the window size needed for competitive performance.

**Table 3.** Effect of window size on QuImputer whole-genome performance.

| Metric | w=25 | w=30 | Change (w=30 - w=25) |
| --- | --- | --- | --- |
| Binary concordance, MAF <1% | 0.999012 | 0.999010 | -0.0002 pp |
| Binary concordance, MAF 1-5% | 0.956730 | 0.956665 | -0.0065 pp |
| Binary concordance, MAF 5-10% | 0.884728 | 0.884655 | -0.0073 pp |
| Binary concordance, MAF 10-50% | 0.697346 | 0.698522 | +0.1176 pp |
| Overall binary concordance | 0.982555 | 0.982571 | +0.0016 pp |
| Pooled dosage R2 | 0.014339 | 0.016068 | +0.001728 |

*Differences are computed at full numerical precision and may therefore differ by one unit in the final digit from the difference of the rounded values shown.*

### 3.4 Diagnostics of local information and parameter sensitivity

We examined local LD support in the first 100,000 SNPs of chromosome 1 under the observed-data condition, using 4,000 non-overlapping windows of 25 SNPs. For each masked true ALT-carrier evaluation at an eligible site, we counted markers in the same window with r^2^ > 0.01. Mean partner counts were 0.000, 0.107, 0.273 and 1.436 across the four MAF bins (Figure 4A; Supplementary Table S11). The corresponding numbers of evaluated carriers were 6,002, 30,832, 33,990 and 81,372. These are carrier-weighted summaries: a site can contribute through several samples. The proportions without an observed ALT-carrier partner were 100.000%, 99.562%, 98.797% and 93.253%. These findings describe limited local support for many examined rare-ALT targets; they do not imply that every quadratic term in their windows is absent.

**Figure 4.**
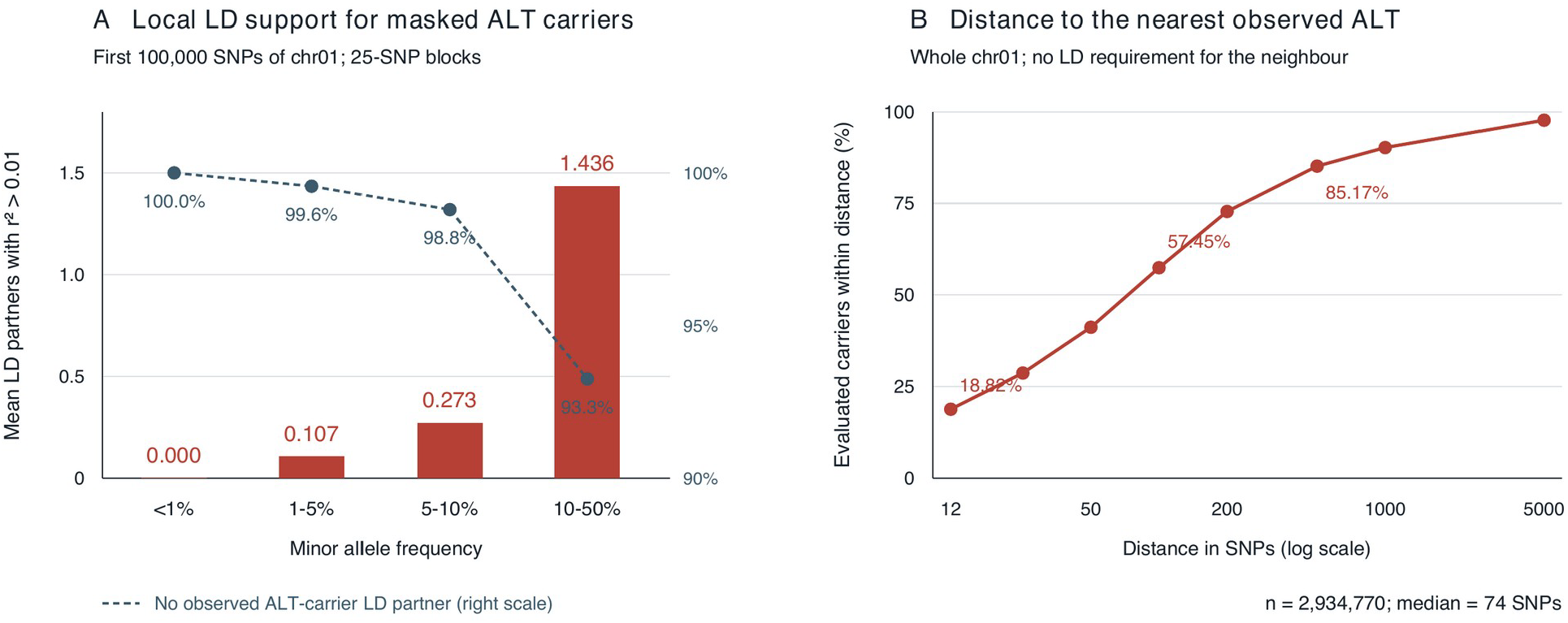
Local LD support and observed ALT spacing on chromosome 1. (A) Mean number of markers with r^2^ > 0.01 in a 25-SNP window for masked true ALT-carrier evaluations in the first 100,000 SNPs. The dashed curve shows the proportion with no observed ALT-carrier LD partner. Bin-specific carrier counts are 6,002, 30,832, 33,990 and 81,372. (B) Cumulative distance to the nearest observed ALT-carrier genotype in the same sample across chromosome 1 (n=2,934,770). Targets span allele frequencies up to 0.50, and neighbour selection does not require LD. The median distance is 74 SNPs (interquartile range 20–227). The distance curve does not measure the capture rate of non-overlapping imputation windows.

A separate analysis across chromosome 1 measured the distance from each eligible masked true ALT-carrier genotype to the nearest observed ALT-carrier genotype in the same sample (n=2,934,770), without restricting targets to rare bins or requiring LD between the two sites. The median distance was 74 SNPs, the interquartile range 20–227 and the 90th percentile 964. Fractions within 12, 100 and 500 SNPs were 18.82%, 57.45% and 85.17% (Figure 4B). This is a descriptive spacing analysis. A distance centred on a target is not the inclusion rule for the non-overlapping windows used in imputation, and an observed ALT at another site need not provide LD support. Observed REF states can also inform a signed-LD model. These distances therefore do not specify a required imputation-window size.

Parameter diagnostics were conducted separately on the 12 archived sparse complete-panel instances with 12-122 variables described in Section 3.6 (Supplementary Table S12). Substituting observed or design-corrected allele frequencies for truth-derived frequencies left the selected solutions at all-REF. In the first 10 instances the observed and corrected AF means were both zero, illustrating the limited effect of reweighting when no ALT evidence is observed. Sweeping w_af/w_ld from 0 to 0.6 produced no true-positive recovery or concordance above all-REF across 1,200 instance-by-ratio combinations. Two instances returned four false ALT calls at ratio zero. At zero prior weight, zero-field variables can have tied optima and the returned solution depends on tie handling. These diagnostics establish sensitivity within the selected instance family; they do not exclude effects of AF estimation, masking or parameterization in the primary whole-genome benchmark.

### 3.5 Exhaustive classical scaling

Exact exhaustive optimization became rapidly more expensive (Table 4; Figure 5). On the same 64-core 100,000-SNP benchmark, runtime increased from 3.0 h at *w* = 20 to 27.9 h at *w* = 25 and 825.7 h at *w* = 30. This establishes an operational boundary for exhaustive enumeration, not for all classical algorithms; simulated annealing, branch-and-bound, and other heuristics were not benchmarked.

**Table 4.** Exhaustive classical runtime on identical hardware (64 cores, 100,000 SNPs).

| Window | Mean active variables | Wall time | Relative to w=20 |
| --- | --- | --- | --- |
| 20 | approximately 16 | 3.0 h | 1.0x |
| 25 | approximately 20 | 27.9 h | 9.3x |
| 30 | approximately 24 | 825.7 h | 274.9x |

*Exact completed wall times were 10,811 s, 100,293 s, and 2,972,437 s for $w=20$ , 25, and 30, respectively. In a separate $w=35$ XC40 stress test, no window completed within a 14-h allocation.*

**Table 5.**
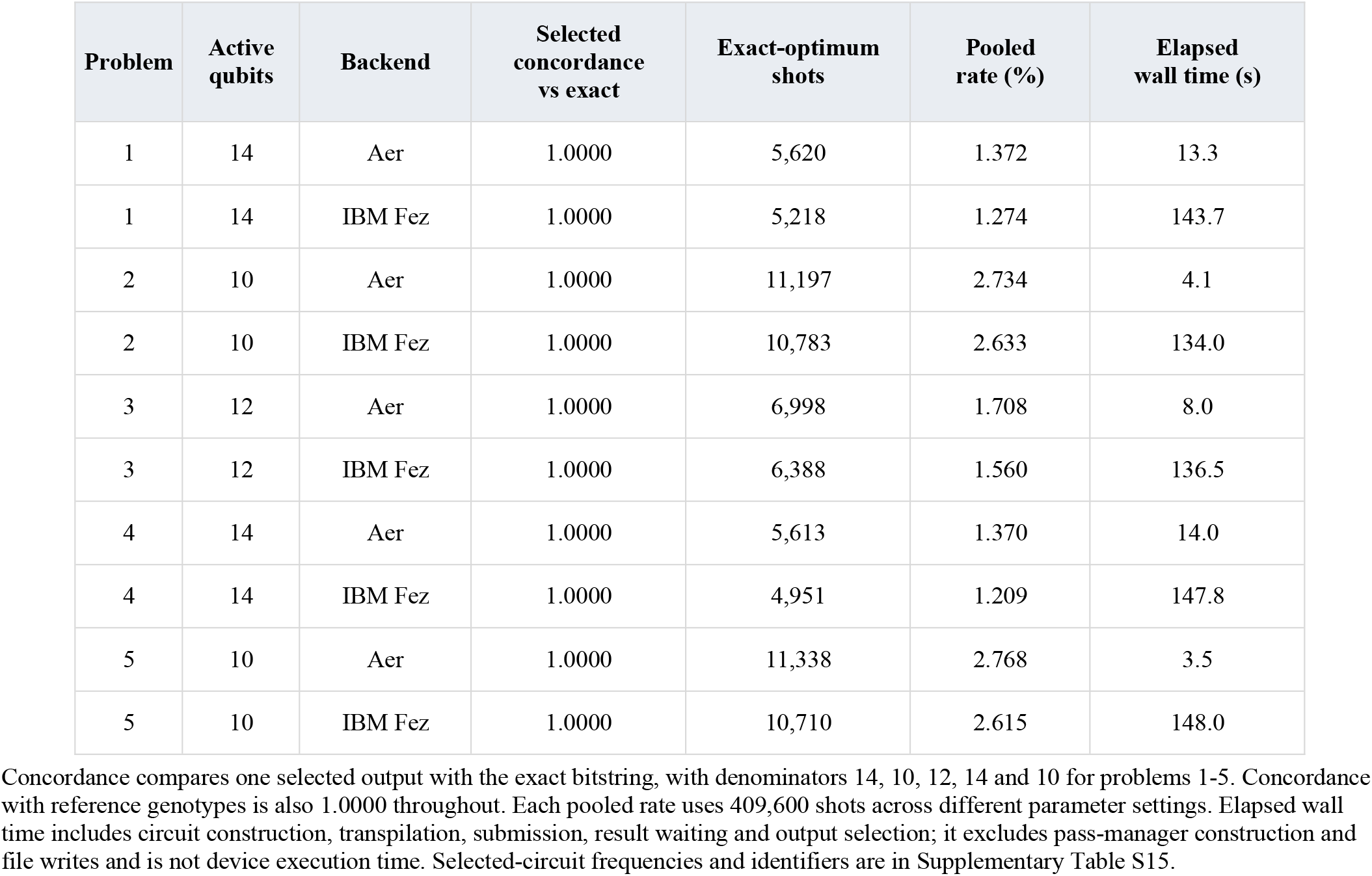
Selected validation problems executed on 18 September 2026. Each row summarizes 100 parameter circuits and 409,600 shots.

| Problem | Active qubits | Backend | Selected concordance vs exact | Exact-optimum shots | Pooled rate (%) | Elapsed wall time (s) |
| --- | --- | --- | --- | --- | --- | --- |
| 1 | 14 | Aer | 1.0000 | 5,620 | 1.372 | 13.3 |
| 1 | 14 | IBM Fez | 1.0000 | 5,218 | 1.274 | 143.7 |
| 2 | 10 | Aer | 1.0000 | 11,197 | 2.734 | 4.1 |
| 2 | 10 | IBM Fez | 1.0000 | 10,783 | 2.633 | 134.0 |
| 3 | 12 | Aer | 1.0000 | 6,998 | 1.708 | 8.0 |
| 3 | 12 | IBM Fez | 1.0000 | 6,388 | 1.560 | 136.5 |
| 4 | 14 | Aer | 1.0000 | 5,613 | 1.370 | 14.0 |
| 4 | 14 | IBM Fez | 1.0000 | 4,951 | 1.209 | 147.8 |
| 5 | 10 | Aer | 1.0000 | 11,338 | 2.768 | 3.5 |
| 5 | 10 | IBM Fez | 1.0000 | 10,710 | 2.615 | 148.0 |

**Figure 5.**
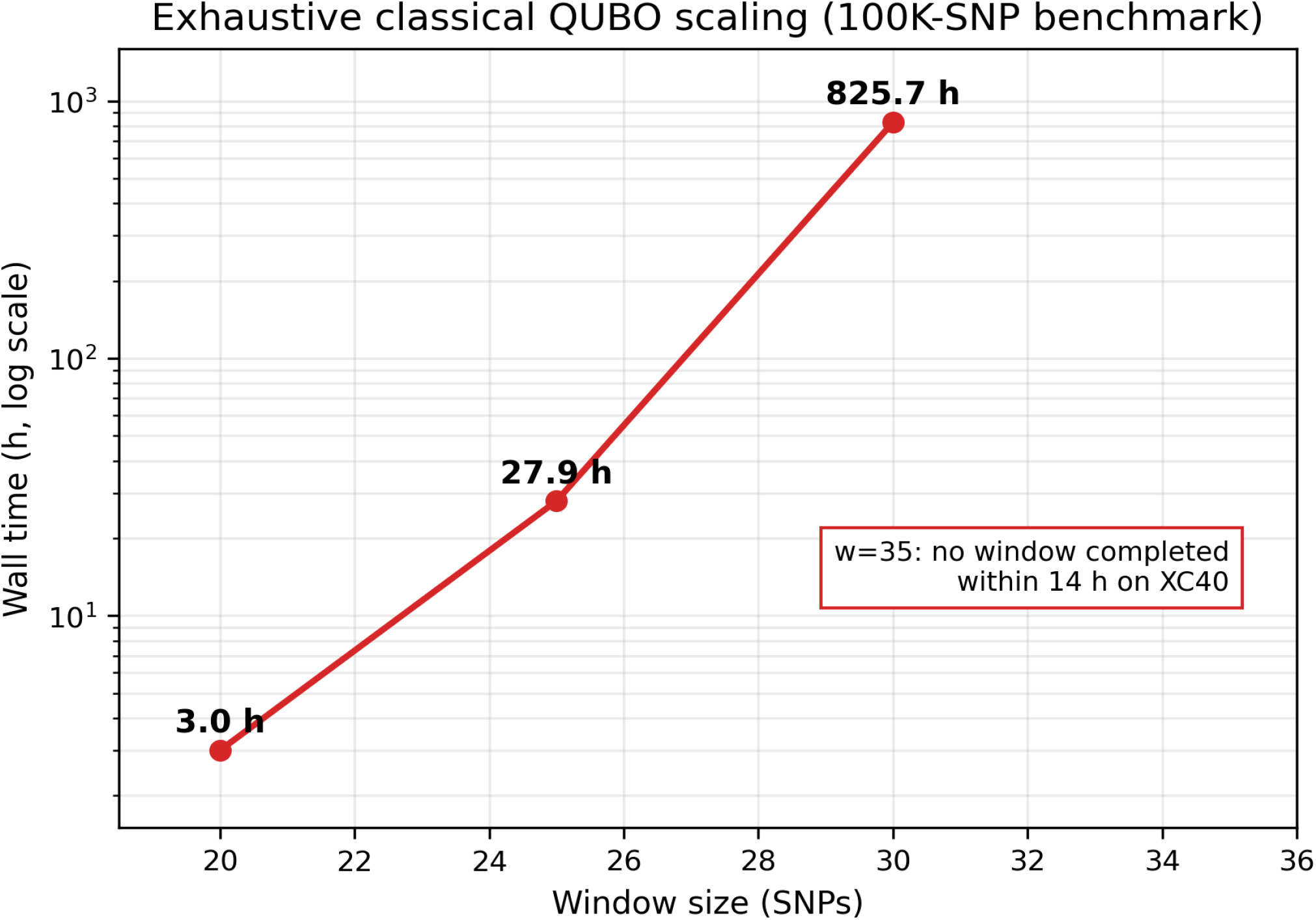
Exhaustive classical QUBO scaling. Wall time is shown on a logarithmic scale. The *w* = 35 entry is a resource-bounded incomplete experiment and is not plotted as a completed runtime.

### 3.6 Quantum-hardware feasibility

Five small problems with 10-14 active qubits were evaluated by exhaustive search, Aer sampling and IBM Fez execution (Table 5; Figure 6). For both backends, selected-output concordance with the exact solution and with the reference genotypes was 1.0000 in every problem. All five exact optima were unique all-REF strings, and the reference genotypes at these masked positions were also all-REF. Four problems had no pairwise couplings; problem 3 had one. The pooled frequency of the exact optimum was 1.370-2.768% for Aer and 1.209-2.633% for IBM Fez. Thus, agreement of the selected output establishes execution and selection of a correct candidate for these simple objectives, without establishing high per-shot success, quantum advantage or state fidelity.

**Figure 6.**
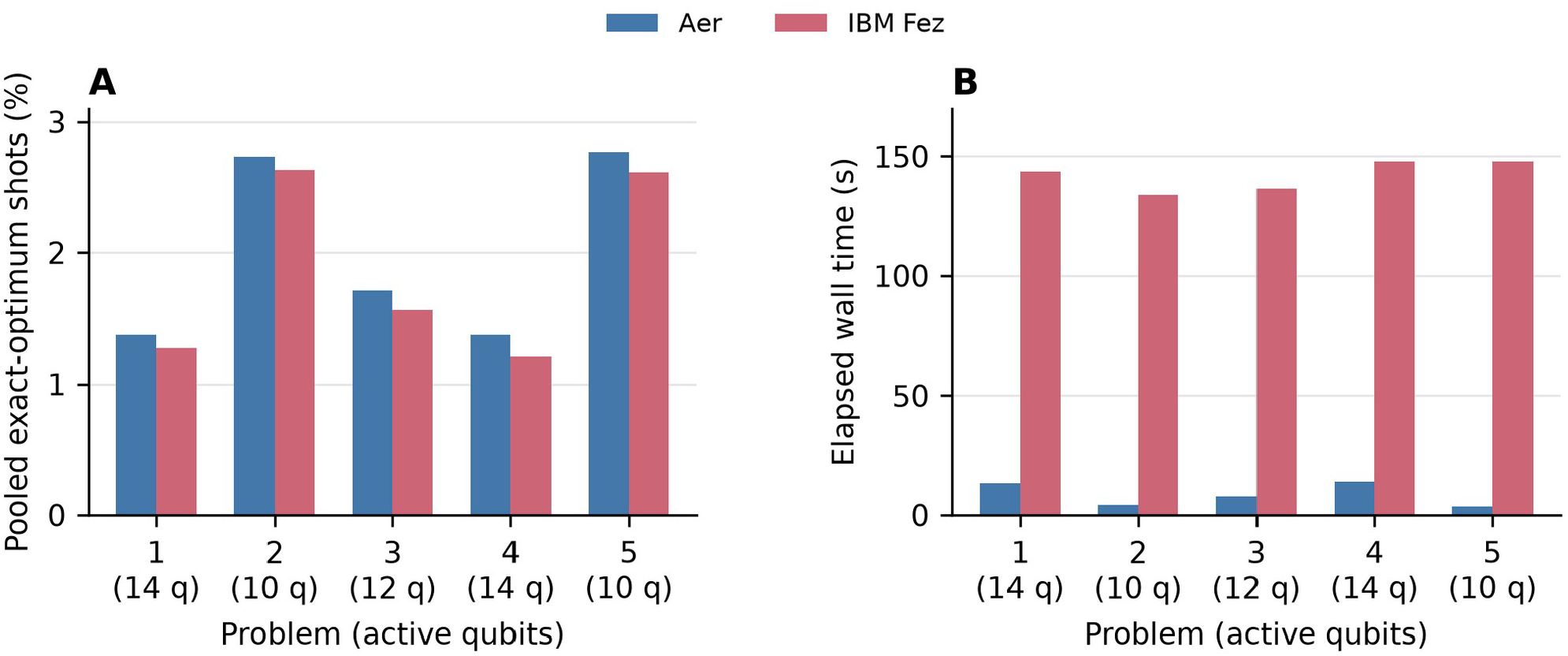
Sampling and elapsed time for the five Table 5 validation problems. (A) Exact-optimum frequency pooled across 100 parameter circuits, each sampled 4,096 times. (B) Elapsed wall time, including classical preparation and output selection as well as waiting for results. Selected-output concordance is 1.0000 for all rows, but the pooled frequencies are much lower. These results concern simple all-REF objectives and do not measure quantum state fidelity.

The October experiment comprised eleven archived sparse objectives with 12-97 variables (A012-A097) and a separate 120-variable objective (B120). All twelve had unique all-REF exact optima. Their coupling graphs contained connected components of at most eight variables, and 75-94% of variables were isolated, allowing exact classical reference calculations (Supplementary Table S13). In the circuit convention z=1-2x, all local fields and retained pairwise coefficients were negative, so all-REF minimized every term simultaneously. Removing pairwise couplings while retaining the local fields gave the same assignment. The archived 122-variable objective used in the parameter diagnostics is distinct from B120 and has no hardware result.

In A012-A049, the selected minimum-energy bitstring equalled the exact optimum in all eight instances (Figure 7; Supplementary Table S10). Across their parameter grids, hardware optimum counts ranged from 5 to 1,285, corresponding to 0.046-0.875 times the noiseless expected counts. No exact optimum was observed for A061, A081 or A097. Their selected outputs matched known biological truth at 58/60, 77/81 and 89/97 variables, respectively, compared with all-REF references of 59/60, 80/81 and 96/97. Missing truth was excluded from these denominators. These shot-resolved results distinguish selection of a correct candidate from its probability of being sampled.

**Figure 7.**
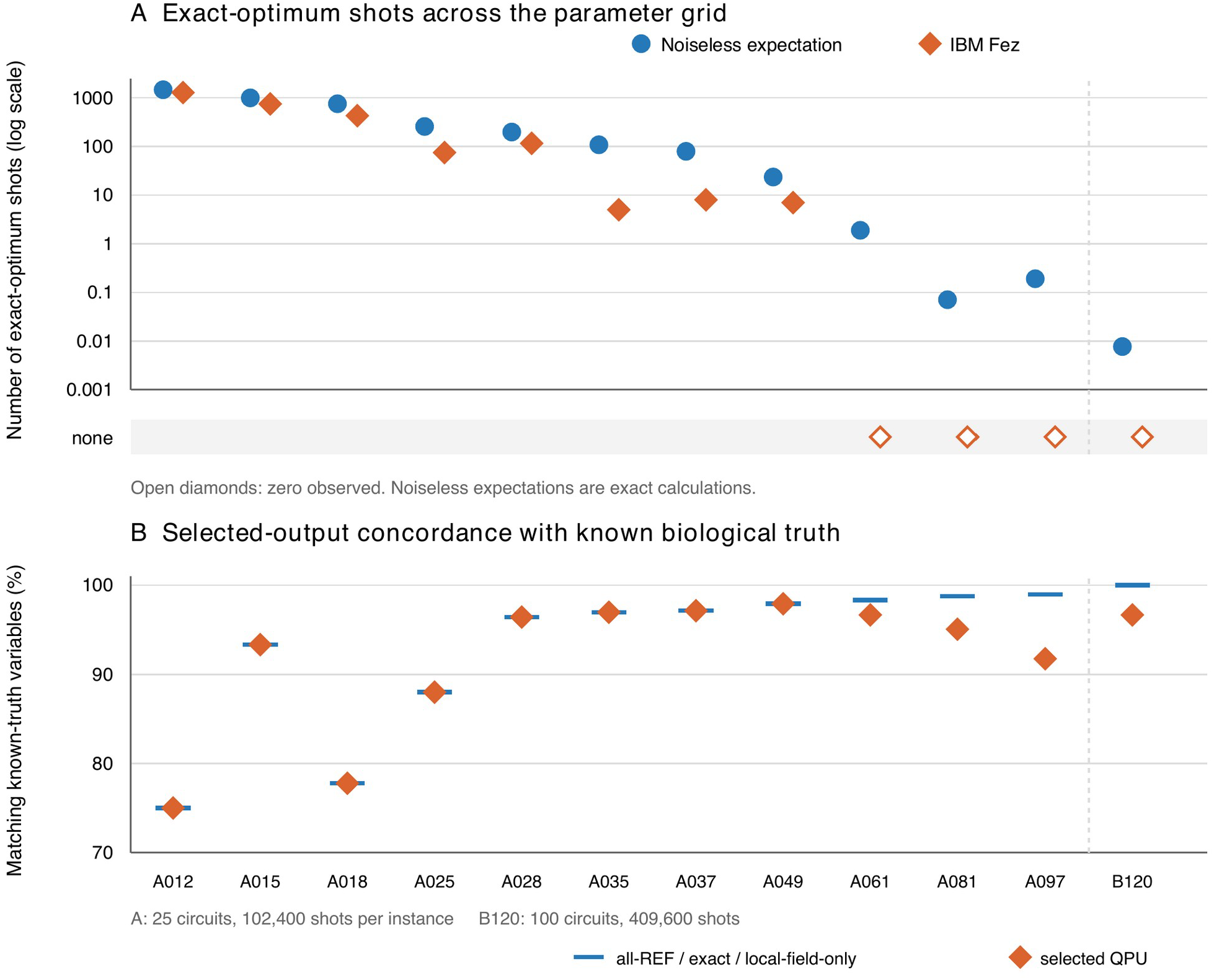
October hardware measurements and exact noiseless references. (A) Exact-optimum shots pooled over the parameter grid: IBM Fez observations and noiseless expectations computed by component-wise simulation. Open diamonds in the separate bottom band indicate zero observations; they are not positive values on the logarithmic axis. (B) Concordance of the selected hardware output with known biological truth, alongside the all-REF, exact-solution and local-field-only references, which coincide. Missing-truth variables are excluded. A instances used 25 circuits and 102,400 shots each; B120 used 100 circuits and 409,600 shots. Labels identify different instances, not a controlled scaling axis. The archived 122-variable instance was not executed and is excluded from this figure.

B120 was executed as a full 120-variable circuit, with 120 active physical qubits, 153 native two-qubit CZ gates and transpiled depth 166. Its 100 parameter circuits returned 409,600 shots and used 110 s of reported QPU time. The selected output matched 116/120 reference states, with an energy 2.4 above the exact minimum; no shot equalled the unique all-REF optimum. The biological truth for this instance was also all-REF, giving a baseline of 120/120. This experiment documents a 120-qubit circuit mapped to 120 active physical qubits, while the objective remains classically tractable by component-wise enumeration.

For A061, A081, A097 and B120, the noiseless expected optimum counts were 1.890, 0.071, 0.191 and 0.00769, respectively. The corresponding probabilities of no optimum in the full finite-shot experiment were 0.1510, 0.9317, 0.8264 and 0.9923. Thus, non-observation of an optimum alone does not diagnose hardware error. Comparisons of the selected outputs and per-variable measurement frequencies provide separate information about deviations from the noiseless model (Supplementary Table S16). Within-instance correlations with binary biological truth, based on ALT shot frequencies and selected bitstrings, are reported in Supplementary Table S5. In B120, all 4,096 strings were distinct within every circuit; this is also expected without noise, with fewer than 0.000121 coinciding shot pairs expected per circuit. Distinct strings alone therefore do not establish uniformity or complete decoherence. These instances differ in genomic content, graph structure and, between A and B, parameter grid and selection settings; they do not isolate a qubit-count or gate-count scaling effect (Preskill, 2018; Gambetta et al., 2017; Harrigan et al., 2021).

## 4. Discussion

QuImputer provides a reduced LD-informed QUBO/Ising formulation for genotype imputation and connects that objective to classical optimization, simulation and quantum-hardware execution. Observed markers enter through local fields, while pairwise terms couple missing variables. This separates the biological objective from the solver used to minimize it and makes the representation available for studies of alternative classical and quantum optimization methods.

The primary observed-data benchmark characterizes the present implementation. Beagle achieved higher sensitivity, F1, MCC and pooled hard-call dosage R^2^, while QuImputer produced fewer false ALT calls. The small genome-wide precision advantage for QuImputer did not extend to gene regions. These results define a conservative calling profile under the evaluated masking regime, and they do not establish a practical advantage over established HMM imputation. Gene-specific examples show heterogeneity but were selected post hoc and do not establish biological or deployment utility.

Local information is one candidate for improving the formulation. The chromosome-1 diagnostic found little within-window LD support for many masked rare-ALT targets. Overlapping windows, observed-marker conditioning beyond the optimization block, or hierarchical summaries of distant LD could change the available information without requiring a single exhaustive search over an entire chromosome. Their effects remain to be measured. The nearest-ALT distance analysis motivates examination of information scope but does not determine a necessary window size or a general limit for local LD methods.

Information conditions must also remain explicit. The secondary complete-panel analyses use genotypes hidden for scoring, and their archived evaluation set differs from the corrected primary set. Their numerical differences from the primary results therefore combine information-source and evaluation-definition changes. We retain them as records of development and sensitivity analyses, without treating those differences as an isolated causal effect of reference information. Parameter selection on an independent observed-data development set and validation under an external reference-panel setting would further characterize generalizability.

The hardware experiments establish a measured execution path from the LD-derived objective to circuits with up to 120 qubits mapped to 120 active physical qubits. The September validation problems and eight October sparse instances through 49 variables returned exact selected outputs. Full shot records for the October series also quantify how often those optima were sampled and permit comparison with noiseless finite-shot expectations. All October optima were all-REF, including in instances containing true ALT carriers, so solving these objectives exactly does not ensure biological accuracy. In the 18-variable instance, for example, the four true ALT positions form a connected component, but the pairwise preference for agreement and REF-favouring fields select all-REF. The small connected components make the present objectives classically tractable. Problems with nontrivial optima and effective classical baselines remain priorities for future solver comparisons.

Several features limit transfer beyond this experiment. The binary encoding cannot emit heterozygous dosage, and the physical-distance decay and disjoint windows simplify genomic structure. The 60% REF/90% ALT masking is a heuristic stress test, while the uniform-masking and multi-seed checks use the secondary condition. Beagle was run with default settings and no genetic map. The accession panel is 84% japonica and 7% indica after selection by missingness, so it is not representative of the full collection. The separate effects of population composition, masking and LD estimation were not measured.

Exhaustive runtime quantifies the cost of the classical solver used here, rather than the limit of all classical optimization. The small connected components in the sparse hardware series demonstrate that raw qubit count can overstate computational difficulty. Comparisons with classical heuristics or other exact methods, together with richer imputation models, are natural extensions of this formulation. The present contribution is the explicit representation and tested execution path on which such comparisons can be built.

## 5. Conclusion

QuImputer encodes allele-frequency priors and signed LD in a reduced QUBO/Ising objective for genotype imputation and supports execution by classical and quantum solvers. We executed selected objectives on IBM hardware using circuits with up to 120 qubits mapped to 120 active physical qubits and retained raw measurements linking each result to its circuit and objective. The primary evaluated configuration achieved lower sensitivity, F1, MCC and pooled hard-call dosage R^2^ than Beagle, with fewer false ALT calls. Diagnostics identified limited local support in the examined chromosome-1 subset, and the hardware objectives had simple all-reference optima. The contribution is an explicit LD-informed formulation with a verified hardware execution path, providing a basis for investigating richer information models and future optimization methods.

## Supporting information

Supplementary data and analysis code

## Data availability

The 3,000 Rice Genomes Project variant resource is publicly available through the European Nucleotide Archive under PRJEB6180 and through Rice SNP-Seek. Supplementary Data 1 identifies the 100 accessions and their quality-control metrics. QuImputer_supplementary_data.zip provides numerical source data for the manuscript tables and figures, together with hardware coefficients, variable mappings, exact references, selected outputs, execution identifiers and settings. Primary, secondary and diagnostic analyses are explicitly labelled. Full September count distributions and October counts and ordered shot arrays, logical and transpiled circuits, software snapshots and noiseless replicate records are retained by the authors and available from the corresponding author on request. Full per-genotype benchmark prediction matrices and complete input VCFs are not included; input hashes and variant/sample mappings document their identity.

## Code availability

A patent application related to QuImputer has been filed. QuImputer_supplementary_data.zip includes a checker for the supplied selected-output energies and concordances and scripts to regenerate Figures 1, 2, 3, 4, 6, 7, S1 and S2. Execution source snapshots, full measurement-verification code and the noiseless-analysis script are available from the corresponding author on request. The complete QuImputer implementation and benchmarking pipeline are being prepared for release in a public repository with a citable DOI at journal publication, subject to institutional software-release and licensing review. Requests for access for peer review will be accommodated through the corresponding author.

## Author contributions

T.-H.L. conceived the study and developed the QuImputer formulation. M.K. and T.-H.L. conducted the computational analyses, interpreted the results, and prepared the manuscript. Both authors reviewed and approved the final version.

## Acknowledgments

The authors acknowledge the IBM Quantum Platform for access to IBM Fez and the Agricultural Biotechnology Supercomputing Center, National Institute of Agricultural Sciences, for Cray XC40 resources.

## Use of generative artificial intelligence

Google Gemini, Anthropic Claude and OpenAI Codex were used under author supervision for language editing, code refactoring, figure preparation and checks of consistency between the manuscript and source data. The authors are responsible for the scientific interpretation and final manuscript.

## Ethics statement

No ethical approval was required. The study used publicly available rice genotype data and did not collect new biological material.

