## Supplementary information for "QuImputer: an LD-informed QUBO/Ising formulation of genotype imputation with quantum-hardware execution"

Tables S1-S16 and Figures S1-S3 support the main manuscript. Primary observed-data results are in S6, S7 and S9 and Figures S1-S2; S2-S4 and S8 are secondary developmental analyses. S5, S10, S13 and S16 describe the October hardware series, S15 the separate September Table 5 experiment, and S12 the archived 12-122-variable diagnostics. S11 reports local-information diagnostics and S14 panel composition. Numerical sources and core hardware records are in QuImputer_supplementary_data.zip. Full measurements, circuits and execution code are retained by the authors and available from the corresponding author on request.

#### Supplementary Table S1 Original chromosome 1 encoding summary

REF and ALT-carrier states map to spins -1 and +1. This original chromosome-1 tabulation uses masked positions with allele-index sums 0, 1 or 2; it is an encoding diagnostic, not the corrected primary evaluation set.

| Metric | Value |
|---|---|
| Tabulated positions with allele-index sum 0, 1 or 2 | 212,600,711 |
| Allele-index sum 0 | 208,841,395 (98.23%) |
| Allele-index sum 1 | 2,050,597 (0.96%) |
| Allele-index sum 2 | 1,708,719 (0.80%) |
| Fraction with allele-index sum 1 | 0.96% |

The 212,600,711 positions exclude missing truth and other index sums. Sum 1 denotes a REF/index-1 pair; sum 2 can also arise from multi-ALT records and is not uniquely homozygous ALT. The 0.96% entry is therefore not a corrected estimate of all heterozygotes. Primary scoring and oracle binary recoding are defined in Methods.

#### Supplementary Table S2. Secondary uniform-masking analysis under the complete-panel condition

This 100,000-SNP chr01 comparison uses allele-frequency-independent 70% masking, complete-panel LD and allele frequencies, and the previous evaluation definition. The sign pattern persists within this secondary condition; it does not establish robustness of the corrected primary benchmark.

| MAF bin | Asymmetric QuImputer | Asymmetric Beagle | Uniform 70% QuImputer | Uniform 70% Beagle |
|---|---|---|---|---|
| <1% | 0.9989 | 0.9990 | 0.9989 | 0.9988 |
| 1-5% | 0.9638 | 0.9620 | 0.9694 | 0.9606 |
| 5-10% | 0.9145 | 0.8982 | 0.9358 | 0.9330 |
| 10-50% | 0.6966 | 0.8449 | 0.7502 | 0.8799 |
| Overall | 0.9801 | 0.9845 | 0.9843 | 0.9874 |

| MAF bin | Asymmetric QuImputer-Beagle | Uniform QuImputer-Beagle | Qualitative sign retained? |
|---|---|---|---|
| 1-5% | +0.18 pp | +0.88 pp | Yes |
| 5-10% | +1.63 pp | +0.28 pp | Yes |
| 10-50% | -14.8 pp | -13.0 pp | Yes (deficit in both) |

#### Supplementary Table S3. Secondary multi-seed analysis under the complete-panel condition

Five masking seeds were evaluated on 100,000 chr01 SNPs with complete-panel statistics and the previous evaluation definition. QuImputer-Beagle concordance differences were positive in both rare bins for all seeds. This documents repeatability within the secondary condition; primary results are in Table S6.

| Seed | QuImputer overall | Beagle overall | QuImputer 1-5% | Beagle 1-5% | QuImputer 5-10% | Beagle 5-10% |
|---|---|---|---|---|---|---|
| 42 | 0.9801 | 0.9845 | 0.9638 | 0.9620 | 0.9145 | 0.8982 |
| 43 | 0.9799 | 0.9843 | 0.9636 | 0.9613 | 0.9132 | 0.8981 |
| 44 | 0.9801 | 0.9845 | 0.9645 | 0.9622 | 0.9149 | 0.9030 |
| 45 | 0.9800 | 0.9843 | 0.9635 | 0.9610 | 0.9131 | 0.9002 |

| Seed | QuImputer overall | Beagle overall | QuImputer 1-5% | Beagle 1-5% | QuImputer 5-10% | Beagle 5-10% |
|---|---|---|---|---|---|---|
| 46 | 0.9800 | 0.9846 | 0.9639 | 0.9614 | 0.9136 | 0.9018 |
| Mean | 0.9800 | 0.9844 | 0.9639 | 0.9616 | 0.9139 | 0.9003 |
| SD | 0.0001 | 0.0001 | 0.0003 | 0.0004 | 0.0007 | 0.0019 |

### Supplementary Table S4. Parameter selection under the complete-panel development condition

The 1,000,000-SNP development grid at w=25 used complete-panel statistics and the previous evaluation definition. Its 30 weight combinations contain 21 distinct ratios. Selected weights w_ld=0.5 and w_af=0.3 were applied unchanged to the primary benchmark. Source: secondary_completepanel/parameter_grid_completepanel.csv.

| $w_{ld}$ / $w_{af}$ | 0.01 | 0.05 | 0.1 | 0.3 | 0.5 | 1.0 |
|---|---|---|---|---|---|---|
| 0.5 | 0.9804 | 0.9808 | 0.9815 | **0.9824** | 0.9822 | 0.9819 |
| 1.0 | 0.9803 | 0.9805 | 0.9808 | 0.9819 | 0.9823 | 0.9822 |
| 2.0 | 0.9803 | 0.9804 | 0.9805 | 0.9813 | 0.9817 | 0.9823 |
| 3.0 | 0.9803 | 0.9803 | 0.9804 | 0.9808 | 0.9814 | 0.9820 |
| 5.0 | 0.9803 | 0.9803 | 0.9804 | 0.9806 | 0.9808 | 0.9815 |

Rows are LD weights and columns are allele-frequency weights; cells are binary concordance. Bold marks the highest reported value, 0.9824 (Figure S3). Generalization requires independent observed-data assessment.

#### Supplementary Table S5 Shot-based binary correlations in October selected circuits

For the minimum-energy selected circuit, per-variable ALT shot frequency is correlated with binary biological truth at known-truth positions. Signed Pearson r, $r^2$ and the separate selected-bitstring $r^2$ are shown. These within-instance statistics do not measure diploid dosage accuracy or probability calibration. Circuit indices are zero-based in parameter-grid order.

| Instance | Known truth n | Selected circuit | Shot Pearson r | Shot $r^2$ | Selected-bit $r^2$ |
|---|---|---|---|---|---|
| A012 | 12 | 2 | 0.0534 | 0.0029 | NA |
| A015 | 15 | 3 | -0.0757 | 0.0057 | NA |
| A018 | 18 | 3 | 0.9786 | 0.9577 | NA |
| A025 | 25 | 3 | 0.8802 | 0.7748 | NA |
| A028 | 28 | 4 | 0.0852 | 0.0073 | NA |
| A035 | 33 | 4 | 0.0798 | 0.0064 | NA |
| A037 | 35 | 4 | 0.0660 | 0.0044 | NA |
| A049 | 48 | 4 | 0.1212 | 0.0147 | NA |
| A061 | 60 | 20 | 0.1039 | 0.0108 | 0.0003 |
| A081 | 81 | 4 | 0.0288 | 0.0008 | 0.0005 |
| A097 | 97 | 4 | 0.0718 | 0.0052 | 0.0008 |
| B120 | 120 | 9 | NA | NA | NA |

A012-A049 selected constant all-REF strings, making selected-bitstring correlations undefined (NA). B120 has constant all-REF truth, so both correlations are undefined. These selected complete-panel instances do not form a controlled scaling experiment.

### Supplementary Table S6. Per-chromosome primary observed-data results

### S6a. Overall binary concordance

| Chromosome | Evaluated N | QuImputer w=25 | QuImputer w=30 | Beagle 5.4 | Difference w=25 (pp) | Difference w=30 (pp) |
|---|---|---|---|---|---|---|
| chr01 | 194,127,592 | 0.985151 | 0.985149 | 0.986689 | -0.1538 | -0.1540 |
| chr02 | 162,850,102 | 0.983806 | 0.983809 | 0.985097 | -0.1291 | -0.1288 |
| chr03 | 158,567,105 | 0.987051 | 0.987047 | 0.987750 | -0.0699 | -0.0703 |
| chr04 | 183,076,613 | 0.983328 | 0.983342 | 0.986095 | -0.2767 | -0.2753 |

| Chromosome | Evaluated N | QuImputer w=25 | QuImputer w=30 | Beagle 5.4 | Difference w=25 (pp) | Difference w=30 (pp) |
|---|---|---|---|---|---|---|
| chr05 | 147,541,791 | 0.986492 | 0.986504 | 0.987113 | -0.0621 | -0.0609 |
| chr06 | 156,293,057 | 0.980378 | 0.980411 | 0.984024 | -0.3646 | -0.3613 |
| chr07 | 148,807,662 | 0.980862 | 0.980902 | 0.984941 | -0.4079 | -0.4039 |
| chr08 | 157,636,280 | 0.982460 | 0.982475 | 0.984530 | -0.2070 | -0.2055 |
| chr09 | 121,743,334 | 0.983316 | 0.983334 | 0.985540 | -0.2224 | -0.2206 |
| chr10 | 127,536,697 | 0.977291 | 0.977299 | 0.979808 | -0.2517 | -0.2509 |
| chr11 | 164,719,282 | 0.976923 | 0.976954 | 0.981446 | -0.4523 | -0.4492 |
| chr12 | 154,618,149 | 0.982370 | 0.982396 | 0.984504 | -0.2134 | -0.2108 |

#### S6b. QuImputer minus Beagle by MAF bin

| Chromosome | 1-5% w=25 | 1-5% w=30 | 5-10% w=25 | 5-10% w=30 | 10-50% w=25 | 10-50% w=30 |
|---|---|---|---|---|---|---|
| chr01 | +0.0816 | +0.0745 | -0.1186 | -0.1453 | -10.2766 | -10.1822 |
| chr02 | +0.0724 | +0.0657 | -0.1110 | -0.1226 | -6.4557 | -6.3829 |
| chr03 | +0.1206 | +0.1148 | +0.0140 | +0.0294 | -7.0762 | -7.0406 |
| chr04 | +0.0794 | +0.0707 | -0.0763 | -0.0888 | -11.3235 | -11.1943 |
| chr05 | +0.0999 | +0.0970 | +0.0008 | -0.0098 | -6.5666 | -6.4064 |
| chr06 | +0.1072 | +0.1004 | -0.1335 | -0.1404 | -12.9606 | -12.8011 |
| chr07 | +0.1006 | +0.0964 | -0.1295 | -0.1344 | -16.3067 | -16.1205 |
| chr08 | +0.1157 | +0.1090 | -0.1458 | -0.1492 | -8.6852 | -8.5842 |
| chr09 | +0.1074 | +0.1020 | +0.0513 | +0.0461 | -10.9078 | -10.7802 |
| chr10 | +0.1208 | +0.1150 | -0.2559 | -0.2687 | -5.9650 | -5.9166 |
| chr11 | +0.0578 | +0.0443 | -0.1023 | -0.1054 | -12.5357 | -12.3793 |
| chr12 | +0.1298 | +0.1273 | -0.0818 | -0.0824 | -9.0121 | -8.8934 |

Low-MAF concordance is reference-dominated and does not imply higher ALT-carrier sensitivity; direct carrier metrics are in Table S7.

### Supplementary Table S7. Rare-ALT confusion counts and gene-level Type II analysis at w=25 and w=30

Gene-region aggregates cover 29,437 overlapping genes with at least three rare SNPs and 50 masked rare-genotype evaluations. The 28,187-gene comparison additionally requires at least 10 true ALT evaluations. Differences within 0.1 percentage points are tied or similar; overlapping genes contribute independently.

#### S7a. Raw confusion counts underpinning the genome-wide and gene-region rates

| Scope | Outcome | QuImputer | Beagle 5.4 |
|---|---|---|---|
| Genome-wide rare-ALT universe (MAF 1-10%, w=25; N=323,536,811) | TP (ALT→ALT) | 93,515 | 438,435 |
| | FN (ALT→REF) | 17,302,693 | 16,957,773 |
| | FP (REF→ALT) | 116,955 | 681,919 |
| | TN (REF→REF) | 306,023,648 | 305,458,684 |
| Annotated gene regions (w=25; 58,390,209 calls) | TP (ALT→ALT) | 24,590 | 96,819 |
| | FN (ALT→REF) | 3,246,551 | 3,174,322 |
| | FP (REF→ALT) | 30,929 | 116,639 |
| | TN (REF→REF) | 55,088,139 | 55,002,429 |

### S7b. Gene-level summary at w=25 and w=30

#### S7b(i). Gene-level comparison by window size

| Window | QuImputer lower Type II | Beagle lower Type II | Tied or similar within 0.1 pp |
|---|---|---|---|
| 25 | 2,395 (8.50%) | 8,503 (30.17%) | 17,289 (61.34%) |
| 30 | 2,603 (9.23%) | 8,385 (29.75%) | 17,199 (61.02%) |

#### S7b(ii). Gene-region error metrics by window size

| Window | QuImputer sensitivity | Beagle sensitivity | QuImputer Type I | Beagle Type I | QuImputer precision | Beagle precision |
|---|---|---|---|---|---|---|
| 25 | 0.75% | 2.96% | 0.056% | 0.212% | 44.29% | 45.36% |
| 30 | 0.87% | 2.96% | 0.073% | 0.212% | 41.53% | 45.36% |

All values use the primary evaluation set. At w=30, QuImputer gene-region sensitivity increases with lower precision and more false ALT calls; its MCC rises from 0.0519 to 0.0535. Beagle precision exceeds QuImputer at both window sizes.

### S7c. Sensitivity of the rare-ALT comparison to the evaluation-set definition

Stage 0 is the previous evaluation; stage 1 excludes missing truth. Stage 2 jointly corrects MAF coding and excludes ALT-major sites. Stage 3 additionally restricts to exactly coded sites and defines the primary rare-ALT set. The alternative retains eligible ALT-major sites and recodes the target as minor-allele carriage. Metrics are QuImputer w=25 / Beagle; stages re-score existing predictions.

| Evaluation set | Evaluated N | Sensitivity (QI / BG) | Precision (QI / BG) | MCC (QI / BG) |
|---|---|---|---|---|
| 0. Previous definition | 361,853,930 | 1.97% / 4.00% | 66.85% / 40.11% | 0.1088 / 0.1138 |
| 1. + missing truth excluded | 343,985,448 | 1.97% / 4.00% | 72.34% / 50.66% | 0.1136 / 0.1311 |
| 2. Corrected MAF and ALT-major sites excluded | 340,333,852 | 0.54% / 2.52% | 43.89% / 38.90% | 0.0439 / 0.0877 |
| 3. + exact coding (primary) | 323,536,811 | 0.54% / 2.52% | 44.43% / 39.13% | 0.0442 / 0.0882 |
| Alt. definition: minor-allele recoding | 323,806,513 | 0.55% / 2.58% | 40.94% / 38.93% | 0.0424 / 0.0889 |

### Supplementary Table S8 Secondary information condition analyses

S8a compares observed and complete-panel statistics on 100,000 chr01 SNPs at w=25 under the earlier evaluation definition. S8b gives historical whole-genome complete-panel results. Complete-panel statistics include genotypes hidden for scoring; S8b is not a matched comparator for corrected primary Tables 1-2 and S6-S7.

### S8a. Chromosome 1 subset comparison

| MAF bin | Observed-data LD | Complete-panel LD | Difference (complete - observed) | Beagle |
|---|---|---|---|---|
| <1% | 0.9990 | 0.9990 | 0.0000 | 0.9990 |
| 1-5% | 0.9637 | 0.9639 | +0.0002 | 0.9620 |
| 5-10% | 0.9028 | 0.9141 | +0.0113 | 0.8982 |
| 10-50% | 0.6129 | 0.6968 | +0.0839 | 0.8449 |
| Overall | 0.9765 | 0.9801 | +0.0036 | 0.9845 |

### S8b Historical whole genome complete panel results

| Historical metric | Complete-panel condition with previous evaluation definition |
|---|---|
| QuImputer-Beagle binary-concordance margin at w=25, MAF 1-5% | +0.40 pp |
| QuImputer-Beagle binary-concordance margin at w=25, MAF 5-10% | +0.90 pp |
| Rare-ALT sensitivity | QuImputer 12.47%; Beagle 4.00% |
| Genes with lower Type II error | QuImputer 70.1%; Beagle 7.2%; tied 22.7% |
| QuImputer-Beagle margin at w=30, MAF 1-5% / 5-10% | +0.39 / +0.88 pp |

S8b retained missing truth and ALT-major sites. Differences from primary results combine information-source, MAF-coding and scoring changes, so they do not isolate an information-source effect. Sparse hardware instances also use the separate complete-panel condition.

### Supplementary Table S9. Per-chromosome dosage $R^2$ and pooled sufficient statistics

| Chromosome | Evaluated N | QuImputer w=25 | QuImputer w=30 | Beagle 5.4 |
|---|---|---|---|---|
| chr01 | 194,127,592 | 0.013221 | 0.014490 | 0.196868 |
| chr02 | 162,850,102 | 0.011307 | 0.012587 | 0.146156 |
| chr03 | 158,567,105 | 0.007673 | 0.008374 | 0.126194 |
| chr04 | 183,076,613 | 0.015405 | 0.016966 | 0.276635 |
| chr05 | 147,541,791 | 0.009448 | 0.010612 | 0.110001 |
| chr06 | 156,293,057 | 0.017861 | 0.020049 | 0.298905 |
| chr07 | 148,807,662 | 0.021730 | 0.024947 | 0.343265 |
| chr08 | 157,636,280 | 0.011324 | 0.012732 | 0.206551 |
| chr09 | 121,743,334 | 0.011861 | 0.013614 | 0.240247 |
| chr10 | 127,536,697 | 0.009126 | 0.010224 | 0.172680 |
| chr11 | 164,719,282 | 0.020948 | 0.023470 | 0.294515 |
| chr12 | 154,618,149 | 0.012772 | 0.014432 | 0.200573 |
| Pooled | 1,877,517,664 | 0.014339 | 0.016068 | 0.231099 |

The primary evaluation definition applies. Files dosage_sufficient_statistics_w25.csv and dosage_sufficient_statistics_w30.csv provide n, Sx, Sy, Sxy, Sxx and Syy by chromosome; reconstructing pooled correlation reproduces these values within 1e-6.

#### Supplementary Table S10 October hardware outputs and optimum sampling

| Instance | Variables | Selected / known truth | all-REF / known truth | Selected / exact | Exact-optimum shots / all shots |
|---|---|---|---|---|---|
| A012 | 12 | 9/12 | 9/12 | 12/12 | 1,285/102,400 |
| A015 | 15 | 14/15 | 14/15 | 15/15 | 751/102,400 |
| A018 | 18 | 14/18 | 14/18 | 18/18 | 430/102,400 |
| A025 | 25 | 22/25 | 22/25 | 25/25 | 75/102,400 |
| A028 | 28 | 27/28 | 27/28 | 28/28 | 116/102,400 |
| A035 | 35 | 32/33 | 32/33 | 35/35 | 5/102,400 |
| A037 | 37 | 34/35 | 34/35 | 37/37 | 8/102,400 |
| A049 | 49 | 47/48 | 47/48 | 49/49 | 7/102,400 |
| A061 | 61 | 58/60 | 59/60 | 60/61 | 0/102,400 |
| A081 | 81 | 77/81 | 80/81 | 78/81 | 0/102,400 |
| A097 | 97 | 89/97 | 96/97 | 90/97 | 0/102,400 |
| B120 | 120 | 116/120 | 120/120 | 116/120 | 0/409,600 |

Executed on ibm_fez on 6 October 2026. A012-A097 match archived G3 coefficients; B120 was newly documented with the retained selection procedure. The lowest-energy observed string is selected over all circuits with the recorded strict tie rule (Methods 2.4). Truth concordance excludes missing truth; exact concordance uses every variable. All exact optima are unique all-REF. A has 25 circuits/102,400 shots per instance; B120 has 100 circuits/409,600 shots. Identifiers and noiseless references are in S16.

A035, A037, A049 and A061 exclude 2, 2, 1 and 1 missing-truth variables, respectively. The September Table 5 problems and archived 122-variable objective are separate. These dated October measurements do not establish identity with historical executables.

### Supplementary Table S11 Local LD support and observed ALT spacing

S11a uses 4,000 non-overlapping 25-SNP windows in the first 100,000 chr01 SNPs and post-masking observed LD. Partners are within-window markers with $r^2 > 0.01$. Means are weighted over masked true ALT-carrier evaluations, with bin denominators 6,002, 30,832, 33,990 and 81,372. REF-partner percentages use masked true REF evaluations separately (Figure 4A).

| MAF bin | Mean LD partners in window | Masked ALT carriers with no observed ALT partner | Masked REF genotypes with no observed REF partner |
|---|---|---|---|
| <1% | 0.0000 | 100.000% | 99.95% |
| 1-5% | 0.1069 | 99.562% | 98.34% |
| 5-10% | 0.2727 | 98.797% | 95.76% |
| 10-50% | 1.4365 | 93.253% | 82.79% |

S11b Distance to the nearest observed ALT carrier across chromosome 1

| Distance threshold in SNPs | Fraction within distance |
|---|---|
| 12 | 18.82% |
| 25 | 28.70% |
| 50 | 41.20% |
| 100 | 57.45% |
| 200 | 72.80% |
| 500 | 85.17% |
| 1000 | 90.23% |
| 5000 | 97.70% |

S11b covers 2,934,770 eligible masked ALT-carrier evaluations across chr01, including common-frequency targets and AF=0.50. Neighbours are in the same sample without an LD requirement. Distance quantiles (SNPs): 5th 3, 10th 5, 25th 20, median 74, 75th 227, 90th 964, 95th 2529. These centred distances do not measure capture by non-overlapping windows or a required window size.

### Supplementary Table S12 Parameter diagnostics on selected sparse instances

Both diagnostics use the 12 archived complete-panel objectives with 12-122 variables, excluding B120. Exact solutions use connected-component enumeration. These selected-instance diagnostics do not exclude AF bias, masking or parameter effects in the primary benchmark.

| Experiment | Manipulation | Result | Interpretation |
|---|---|---|---|
| AF substitution | Truth-derived, observed and design-corrected AF on 12 selected instances | All three sources returned all-REF; 10 instances had zero observed and corrected mean AF | Sensitivity within this selected family |
| Weight-ratio sweep | 100 ratios from 0 to 0.6 on each of 12 instances | No true positives or concordance above all-REF; at ratio zero, two instances returned four FP calls | Zero-prior ties and instance selection limit generalization |
| Weight scaling | Common positive scaling of both weights | Exact argmin unchanged; 21 distinct ratios in the 30-point development grid | The exact objective depends on the weight ratio |

**Supplementary Table S13 Exact references and coupling-graph structure**

A objectives match archived G3 coefficients; B120 is newly documented, and archived A122 was not executed on hardware. Component enumeration gives a unique all-REF optimum for each objective. Removing pairwise J while retaining h gives the same assignment. Circuit convention z=1-2x reverses local-field signs relative to s=2x-1, preserving pairwise coefficients.

| Instance | Variables | Pairwise terms | Largest component | Isolated variables | Exact assignment | Number of optima |
|---|---|---|---|---|---|---|
| A012 | 12 | 3 | 3 | 9 | all-REF | 1 |
| A015 | 15 | 3 | 3 | 12 | all-REF | 1 |
| A018 | 18 | 6 | 4 | 14 | all-REF | 1 |
| A025 | 25 | 3 | 3 | 22 | all-REF | 1 |
| A028 | 28 | 3 | 3 | 25 | all-REF | 1 |
| A035 | 35 | 6 | 4 | 31 | all-REF | 1 |
| A037 | 37 | 6 | 4 | 33 | all-REF | 1 |
| A049 | 49 | 3 | 3 | 46 | all-REF | 1 |
| A061 | 61 | 6 | 4 | 57 | all-REF | 1 |
| A081 | 81 | 29 | 8 | 71 | all-REF | 1 |
| A097 | 97 | 28 | 8 | 89 | all-REF | 1 |
| B120 | 120 | 9 | 4 | 113 | all-REF | 1 |
| A122* | 122 | 62 | 12 | 108 | all-REF | 1 |

The October objectives have components of at most eight variables and 75-94% isolated variables; A122 has a largest component of 12. A plus B120 form the October series, whereas A plus A122 underlie S12. Full coefficients, energies, exact strings and genomic mappings are in the supplementary data: core/October_instances.json and G3_instances_hJ.json under quantum_hardware/. Hardware outputs and truth denominators are in S10.

### Supplementary Table S14. Variety-group composition of the 100-accession panel

Neighbour-Joining variety-group assignments from the 3,000 Rice Genomes metadata were joined by SRA accession; all 100 matched. Source labels are unchanged. Selection used sample missingness, not variety group.

| Variety group | Panel (n = 100) | 3,000 Rice Genomes collection | Enrichment |
|---|---|---|---|
| japonica groups | 84 (84.0%) | 839 (27.97%) | 3.00 |
| Indica | 7 (7.0%) | 1743 (58.10%) | 0.12 |
| Aus groups | 1 (1.0%) | 215 (7.17%) | 0.14 |
| Aromatic groups | 0 (0.0%) | 68 (2.27%) | absent from panel |
| Intermediate type | 8 (8.0%) | 135 (4.50%) | 1.78 |

Per-accession assignments are in Supplementary Data 1. The panel has threefold japonica enrichment, eightfold indica depletion and no aromatic accessions.

### Supplementary Table S15 September execution records for main Table 5

The five 18 September 2026 problems come from the first 20-SNP chr01 window, selected after loading 50,000 SNPs. Complete-panel statistics used LD weight 0.5, AF weight 0.3, distance decay 10,000 bp and LD threshold 0.01. These checks are separate from the primary observed-data evaluation.

#### S15a Instance identities and exact references

| Problem | Sample accession | Variables | Pairwise terms | Unique exact bitstring | Exact energy |
|---|---|---|---|---|---|
| 1 | ERS468888 | 14 | 0 | 00000000000000 | -4.200561 |
| 2 | ERS470504 | 10 | 0 | 0000000000 | -3.000561 |
| 3 | ERS468864 | 12 | 1 | 000000000000 | -3.594561 |
| 4 | ERS468482 | 14 | 0 | 00000000000000 | -4.200561 |
| 5 | ERS468827 | 10 | 0 | 0000000000 | -3.000561 |

Bits follow logical variable order (0=REF, 1=ALT carriage). Both backends selected the listed exact strings; reference truth is also all-REF. Variable/SNP order and positions are in problems/problem_map.csv and the public core/September_records.json extracts. Only problem 3 has a pairwise coupling.

Q is upper triangular with each off-diagonal term counted once and the offset included. Circuit spins z=1-2b reverse the local-field signs relative to s=2b-1. QUBO and circuit-Ising energies agree for all five objectives; full-precision values are in T5_instance_coefficients.json.

#### S15b Selected circuits and optimum frequencies

| Problem | Backend | Selected index | Gamma | Beta | Exact shots in selected circuit | Circuits with any exact hit |
|---|---|---|---|---|---|---|
| 1 | Aer | 5 | -1.5708 | 0.0873 | 1/4,096 | 41/100 |
| 1 | IBM Fez | 5 | -1.5708 | 0.0873 | 6/4,096 | 46/100 |
| 2 | Aer | 3 | -1.5708 | -0.2618 | 1/4,096 | 66/100 |
| 2 | IBM Fez | 4 | -1.5708 | -0.0873 | 5/4,096 | 68/100 |
| 3 | Aer | 5 | -1.5708 | 0.0873 | 5/4,096 | 54/100 |
| 3 | IBM Fez | 5 | -1.5708 | 0.0873 | 6/4,096 | 53/100 |
| 4 | Aer | 6 | -1.5708 | 0.2618 | 40/4,096 | 46/100 |
| 4 | IBM Fez | 6 | -1.5708 | 0.2618 | 40/4,096 | 42/100 |
| 5 | Aer | 4 | -1.5708 | -0.0873 | 1/4,096 | 64/100 |
| 5 | IBM Fez | 5 | -1.5708 | 0.0873 | 17/4,096 | 71/100 |

Indices are zero-based: gamma is the outer loop, beta the inner. Each grid has 10 equally spaced values on [-π/2, π/2] and [-π/4, π/4], with 4,096 shots per circuit. Strict minimum-energy selection retains the first candidate on floating-point ties, using saved counts order. It selects the exact energies in S15a, not necessarily the circuit with the most optimum hits. Pooled counts/rates are in main Table 5.

#### S15c Execution identifiers

| Problem | Aer local job UUID | IBM Quantum job ID |
|---|---|---|
| 1 | b6d2ab51-1339-42ce-9388-69b5786d41ef | damdpclr85ps73fcmj80 |
| 2 | 4aea3165-f4ad-4cf6-81e3-3108547bcc7e | damdri82fm4c73f2s3og |
| 3 | ab23ab74-9a25-45cc-9f36-3d4604af7a5b | damdssdr85ps73fcmr5g |
| 4 | c1822a21-dc6a-4c09-9d80-692b43dc6220 | damdu7n8gn2s739l9rm0 |
| 5 | dbfb864a-9146-4f36-b73f-f85fd6136a7d | damdvotr85ps73fcn11g |

Aer IDs are local UUIDs; IBM IDs identify QPU jobs and link the saved QPY circuits. IBM Fez is a 156-qubit Heron processor. Both backends used optimization level 1 without explicit sampler/transpiler seeds. Versions: Python 3.13.12, Qiskit 2.3.1, Aer 0.17.2, IBM Runtime 0.46.1, NumPy 2.4.4. The retained full driver/source snapshot includes uncommitted changes relative to its recorded Git HEAD.

All 10 complete execution records and 1,000 circuit count distributions, with the offline verifier, are retained and available from the corresponding author on request. The public core records support selected-string energy and concordance checks; replaying selection and recounting shots require the full records. Exact references were used only for evaluation/recording. Table 5 wall time includes preparation, submission, waiting and selection, excludes pass-manager construction and file writes, and is not pure device time.

#### Supplementary Table S16 October execution records and noiseless comparisons

One Sampler job per instance used optimization level 0, transpiler seed 42, 4,096 shots per circuit and disabled twirling/dynamical decoupling. Each variable mapped to one active physical qubit on 156-qubit ibm_fez. IBM-reported QPU usage totalled 431 s on an Open-plan account, distinct from wall time. Versions: Python 3.13.12, Qiskit 2.3.1, IBM Runtime 0.46.1, NumPy 2.4.4. Core execution metadata are in core/October_execution.json; full source/backend snapshots are available from the corresponding author on request.

#### S16a Instance identities and IBM job identifiers

| Instance | Sample accession | Window start SNP index | IBM Quantum job ID |
|---|---|---|---|
| A012 | ERS470256 | 18 | db2bmpu8v0ts73c2j4h0 |
| A015 | ERS470669 | 23 | db2bnv7r11fs7396gih0 |
| A018 | ERS470256 | 31 | db2bornr11fs7396gk9g |
| A025 | ERS470334 | 0 | db2bpmvr11fs7396glm0 |
| A028 | ERS470222 | 0 | db2bqc7r11fs7396gmag |
| A035 | ERS470448 | 0 | db2br77r11fs7396gn60 |
| A037 | ERS470448 | 0 | db2bs7vr11fs7396go70 |
| A049 | ERS470463 | 0 | db2bspc7f06c73apjma0 |
| A061 | ERS470463 | 0 | db2btak2ljfc73d489ig |
| A081 | ERS470222 | 0 | db2bu0vr11fs7396gq4g |
| A097 | ERS468339 | 0 | db2buic2ljfc73d48asg |
| B120 | ERS468324 | 0 | db2bv7k2ljfc73d48bi0 |

Window starts are zero-based chr01 SNP indices. A used 100,000 SNPs, windows int(target/0.75)+5, and LD $r^2$ thresholds 0.01 (12/15 variables), 0.1 (18) or 0.3 (25-97); selection required three or more couplings and mixed binary truth. One coefficient/truth match was found per A instance. B120 used 50,000 SNPs, 200-SNP windows, $r^2$=0.5 and the first eligible 120-variable problem; coefficient identity with the March job is unconfirmed. Input VCF hashes are retained.

#### S16b Circuit resources and exact noiseless references

| Instance | Native CZ gates | ISA depth | QPU seconds | Expected optimum shots | P(no optimum) | Mean absolute ALT-freq. error |
|---|---|---|---|---|---|---|
| A012 | 27 | 80 | 29 | 1469.2 | $<10^{-6}$ | 0.0264 |
| A015 | 33 | 92 | 29 | 998.74 | $<10^{-6}$ | 0.0263 |
| A018 | 54 | 124 | 29 | 756.84 | $<10^{-6}$ | 0.0215 |
| A025 | 66 | 108 | 29 | 257.9 | $<10^{-6}$ | 0.0246 |
| A028 | 27 | 92 | 29 | 198.22 | $<10^{-6}$ | 0.0141 |
| A035 | 42 | 127 | 29 | 108.4 | $<10^{-6}$ | 0.0251 |
| A037 | 42 | 127 | 29 | 79.526 | $<10^{-6}$ | 0.0246 |
| A049 | 33 | 92 | 29 | 23.542 | $<10^{-6}$ | 0.0168 |
| A061 | 78 | 161 | 29 | 1.8902 | 0.1510 | 0.0201 |
| A081 | 370 | 415 | 30 | 0.070709 | 0.9317 | 0.0408 |
| A097 | 299 | 381 | 30 | 0.19064 | 0.8264 | 0.0360 |
| B120 | 153 | 166 | 110 | 0.0076856 | 0.9923 | 0.0266 |

Expected hits are the sum of 4,096 times each circuit optimum probability pc. P(no optimum) is the product of (1-pc)^4,096 over the grid for independent noiseless shots; it does not describe the selected output truth score. ALT-frequency error averages absolute hardware-minus-noiseless differences across variables and circuits. Calculations use exact components, without a full 120-qubit statevector.

#### S16c Finite-shot distribution of the selected output

| Instance | Hardware truth matches | Noiseless median | Replicates at or below hardware | Noiseless distribution (matches: number of replicates) |
|---|---|---|---|---|
| A061 | 58/60 | 59/60 | 43/500 | 58: 43, 59: 433, 60: 24 |
| A081 | 77/81 | 79/81 | 22/500 | 77: 22, 78: 196, 79: 191, 80: 83, 81: 8 |
| A097 | 89/97 | 95/97 | 0/500 | 94: 20, 95: 332, 96: 126, 97: 22 |
| B120 | 116/120 | 117/120 | 22/500 | 116: 22, 117: 242, 118: 179, 119: 55, 120: 2 |

Each instance has 500 independent finite-shot replicates from exact component probabilities, resetting seed 42 and matching the hardware grid, shots, accumulation order and strict first-observation tie rule. Counts are descriptive, not exact or multiplicity-adjusted tail probabilities; 0/500 does not imply zero probability. A012-A049 selected all-REF in every replicate. All replicate strings, energies and circuit indices are retained in noiseless_recomputed.json and available from the corresponding author on request.

B120 returned 4,096 distinct strings in every circuit. The noiseless expected number of coinciding pairs is at most 0.00012075 per circuit (0.00048478 over the grid), so distinctness does not establish uniformity or complete decoherence. Hardware ALT frequencies differed by more than three ideal standard errors in 31.675% of variable-circuit pairs; this is descriptive, not a failed-qubit fraction or global test probability.

### Supplementary figures

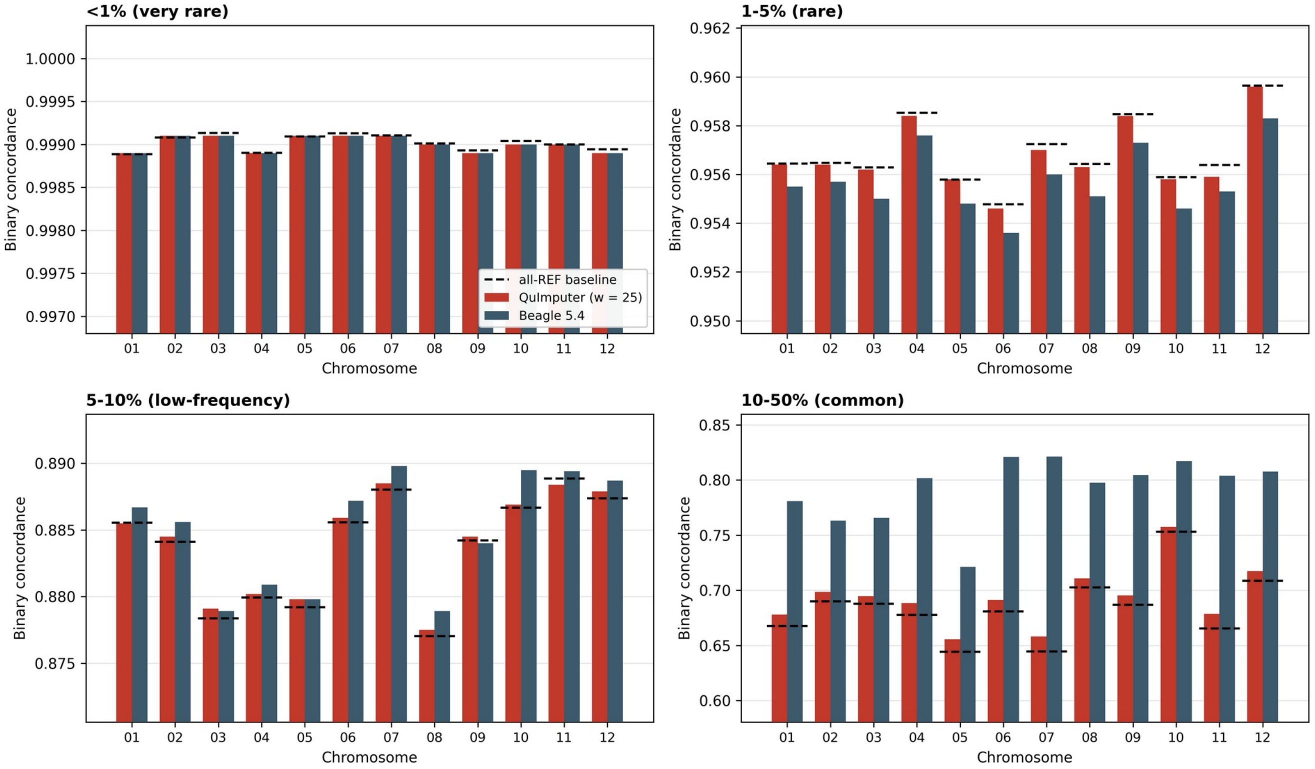


Supplementary Figure S1. Per-chromosome concordance under the corrected primary observed-data definition at w=25. Panels show the four MAF bins, with all-REF baseline segments for each chromosome. Low-MAF concordance must be interpreted alongside direct ALT-carrier metrics.

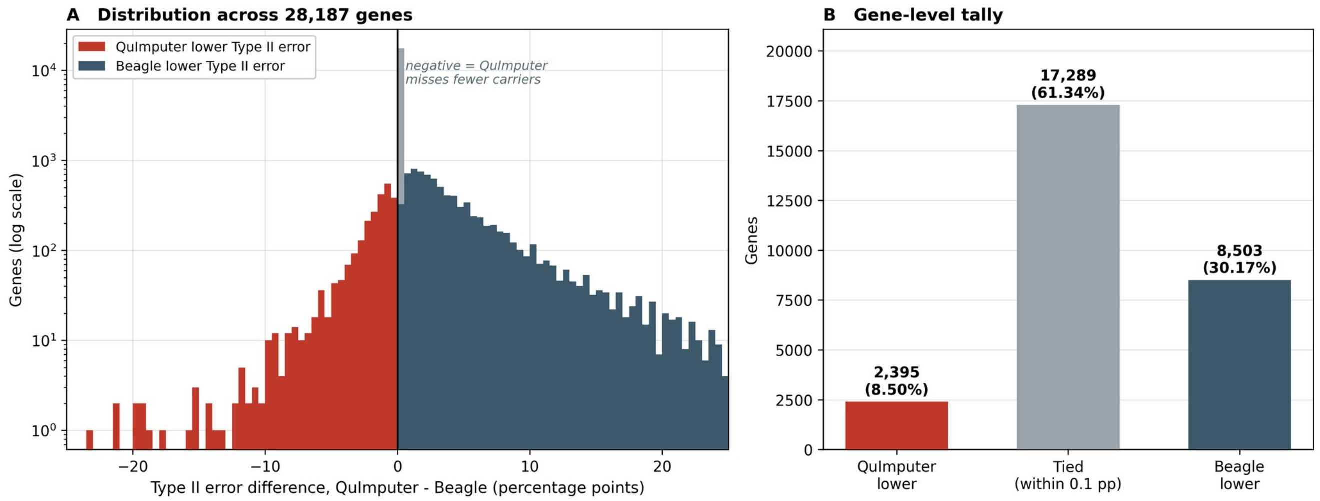


Supplementary Figure S2. Per-gene Type II error differences across 28,187 eligible genes at w=25. (A) QuImputer minus Beagle, in percentage points, on a logarithmic count axis. Negative values favour QuImputer. (B) Counts using a ±0.1 percentage-point tolerance for tied or similar performance: QuImputer lower in 2,395 genes, Beagle lower in 8,503, and 17,289 tied or similar.

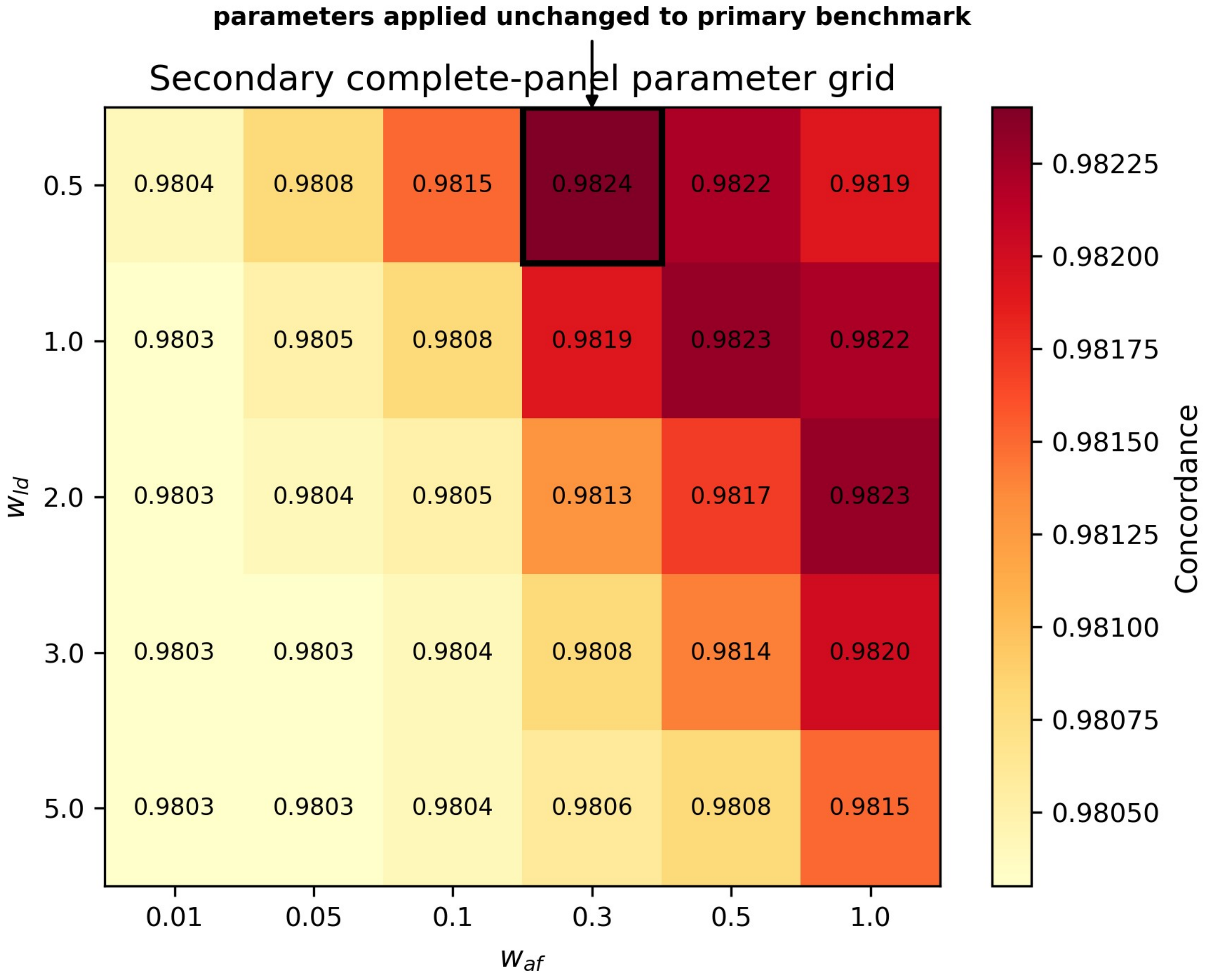


Supplementary Figure S3. Parameter grid under the secondary complete-panel development condition and previous evaluation definition. The selected weights w_ld=0.5 and w_af=0.3 were applied unchanged to the primary benchmark. The grid documents development rather than primary performance.
