## Supplementary data and analysis code for "QuImputer: an LD-informed QUBO/Ising formulation of genotype imputation with quantum-hardware execution": check_core_summaries.py

```
"""Check supplied selected-output summaries; does not replay all measurements."""
from pathlib import Path
import json, math, sys

root=Path(sys.argv[1]) if len(sys.argv)>1 else Path(__file__).resolve().parents[2]
core=root/'quantum_hardware/core'
read=lambda p:json.loads(p.read_text())
def near(a,b):return math.isclose(float(a),float(b),rel_tol=0,abs_tol=1e-8)
def qubo(bits,Q,offset):
    return sum(bits[i]*v*bits[j] for i,row in enumerate(Q) for j,v in enumerate(row))+offset
def ising(bits,h,J):
    z=[1-2*x for x in bits]
    return sum(a*b for a,b in zip(z,h))+sum(float(v)*z[int(k.split(',')[0])]*z[int(k.split(',')[1])] for k,v in J.items())
reports=[]
for r in read(core/'September_records.json')['records']:
    bits=r['final_selection']['bits_logical'];truth=r['truth_bits'];exact=r['exact_bits'];n=r['n_qubits']
    checks={'bit_width':len(bits)==len(truth)==len(exact)==n,
        'selected_Q_energy':near(qubo(bits,r['Q_upper_triangular'],r['offset']),r['final_selection']['energy']),
        'selected_Ising_energy':near(ising(bits,r['h'],r['J']),r['final_selection']['energy']),
        'exact_Q_energy':near(qubo(exact,r['Q_upper_triangular'],r['offset']),r['exact_energy']),
        'truth_concordance':near(sum(a==b for a,b in zip(bits,truth))/n,r['conc_truth']),
        'exact_concordance':near(sum(a==b for a,b in zip(bits,exact))/n,r['conc_exact'])}
    reports.append({'record':r['execution']['backend']+'/'+str(r['table5_problem']),'checks':checks,'pass':all(checks.values())})
instances=read(core/'October_instances.json')['problems'];execution=read(core/'October_execution.json')
for m in read(root/'quantum_hardware/QPU_20261006/paper_metrics.json')['problems']:
    p=instances[m['problem_id']]['problem'];bits=m['selected_bits'];exact=p['exact']['bits'];mask=p['truth_valid_mask'];truth=p['truth_binary_retained_rule']
    checks={'bit_width':len(bits)==p['n_variables']==m['n'],
        'selected_Ising_energy':near(ising(bits,p['h_c'],p['J_c']),m['selected_energy']),
        'selected_Q_energy':near(qubo(bits,p['Q_upper_triangular'],p['offset']),m['selected_energy']+p['C']),
        'exact_energy':near(ising(exact,p['h_c'],p['J_c']),m['exact_energy']),
        'truth_matches':sum(a==b for a,b,v in zip(bits,truth,mask) if v)==m['truth_matches'],
        'truth_denominator':sum(mask)==m['truth_denominator'],
        'exact_matches':sum(a==b for a,b in zip(bits,exact))==m['exact_matches'],
        'job_mapping':execution['job_problem_map']['jobs'][m['job_id']]['problem_id']==m['problem_id']}
    reports.append({'record':m['problem_id'],'checks':checks,'pass':all(checks.values())})
result={'scope':'Selected-string energy/scoring and identity consistency only; raw-count recounting and output reselection require author-retained full records.',
    'all_pass':all(r['pass'] for r in reports),'records':len(reports),'reports':reports}
print(json.dumps(result,indent=2))
sys.exit(0 if result['all_pass'] else 1)
```
