## Supplementary data and analysis code for "QuImputer: an LD-informed QUBO/Ising formulation of genotype imputation with quantum-hardware execution": README_core.txt

Core hardware records accompanying QuImputer, 2026-10-07 v4

September_records.json contains selected fields from all five Aer and five QPU
execution records, including coefficients, coding, variable identities, grids,
selection rules, selected strings, exact references, energies and reported shot
totals. October_instances.json contains all twelve complete problem objects;
October_execution.json contains execution settings, grids, circuit resources,
job identifiers, input hashes and source hashes. October selected outputs and
published metrics are in ../QPU_20261006/paper_metrics.json. Archived G3 coefficients
are in ../G3_instances_hJ.json; the archived 122-variable objective has no QPU result.

Run with Python 3, without IBM access or third-party libraries:
python quantum_hardware/core/check_core_summaries.py .

The checker recomputes energies and concordances of the supplied selected strings,
compares exact-reference energies and checks job/instance identities. It does not
reselect outputs from all measurements or independently recount optimum shots.
Those checks require the full retained counts and ordered shot arrays, which,
together with QPY circuits and execution source snapshots, are available from the
corresponding author on request. File hashes identify the retained source files;
they are not independent proof of their contents. No scientific values were changed
in preparing these extracts.
