## Supplementary data and analysis code for "QuImputer: an LD-informed QUBO/Ising formulation of genotype imputation with quantum-hardware execution": README_source_data.txt

QuImputer supplementary data 2026-10-07 v4

This is the single public data archive accompanying the manuscript and
Supplementary Information. It is self-contained; no second public ZIP is required.
SOURCE_DATA_INDEX.csv maps files to manuscript tables, figures and Methods.

primary_observed/ contains the primary observed-data benchmark summaries.
secondary_completepanel/ contains labelled developmental analyses using the
previous evaluation definition. mechanism/ and runtime/ contain diagnostic and
timing results. SupplementaryData1_accessions.csv identifies the 100 accessions.
Table and figure numbers and numerical results are unchanged from v3.

quantum_hardware/core/ adds coefficients, variable/SNP mappings, selected outputs,
exact references, execution identifiers and settings for September and October.
Read README_core.txt for the precise verification scope. Full counts, ordered
October shots, QPY circuits, software snapshots, full offline verification code
and finite-shot replicate records are retained by the authors and available
through the corresponding author on request; they are not attached in this ZIP.
Full input VCFs and full per-genotype benchmark predictions are not included.

From QuImputer_data/, check supplied hardware summaries with Python 3:
python quantum_hardware/core/check_core_summaries.py .

Regenerate manuscript figures (NumPy, Matplotlib, ReportLab and Poppler):
python scripts/make_archive_figures.py .
python scripts/make_figure6.py .
python scripts/make_figure7_20261006.py quantum_hardware/QPU_20261006/paper_metrics.json /tmp/Figure7.pdf
python scripts/make_figure1_20261007.py /tmp/Figure1.pdf
The first command produces Figures 2, 3, 4, S1 and S2; Figures 5 and S3 remain
embedded in the manuscript files. Set QUIMPUTER_PDFTOPPM or QUIMPUTER_FONT_DIR
if needed for the local Poppler executable and Arial/DejaVu fonts. Use a working
copy for plotting. SOURCE_DATA_SHA256SUMS.txt covers this archive.
