## Supplementary data and analysis code for "QuImputer: an LD-informed QUBO/Ising formulation of genotype imputation with quantum-hardware execution": make_archive_figures.py

```
#!/usr/bin/env python3
# -*- coding: utf-8 -*-
"""Regenerate Figures 2, 3, 4, S1 and S2 from this archive.

Figures 3, S1 and S2 use NumPy and Matplotlib. Figures 2 and 4 are generated
by make_revised_figures.py using ReportLab and Poppler pdftoppm.
Usage: python scripts/make_archive_figures.py [archive_root]
"""
import csv, json, os, sys
import numpy as np
import matplotlib
matplotlib.use('Agg')
import matplotlib.pyplot as plt

ROOT = sys.argv[1] if len(sys.argv) > 1 else os.path.dirname(os.path.dirname(os.path.abspath(__file__)))
OUTD = os.path.join(ROOT, 'figures')
os.makedirs(OUTD, exist_ok=True)
PO = os.path.join(ROOT, 'primary_observed') + os.sep
MECH = os.path.join(ROOT, 'mechanism') + os.sep
QH = os.path.join(ROOT, 'quantum_hardware') + os.sep
RED, SLATE, GREY, AMB = '#C0392B', '#3D5A6C', '#9AA5AB', '#B8860B'


t1 = {r['bin']: r for r in csv.DictReader(open(PO + 'G2_table1.csv'))}
t2 = {(r['scope'], r['method']): r for r in csv.DictReader(open(PO + 'G2_table2.csv'))}


BINS = ['MAF<1%', 'MAF1-5%', 'MAF5-10%', 'MAF10-50%']
LAB = ['<1%\n(very rare)', '1–5%\n(rare)', '5–10%\n(low-frequency)', '10–50%\n(common)']

# =============== FIGURE 3 : rare-ALT error profile =======================
fig, (b1, b2) = plt.subplots(1, 2, figsize=(12.6, 4.9))
met = [('type1', 'Type I error\n(false ALT rate)', 100),
       ('sens', 'Sensitivity\n(true ALT detected)', 100),
       ('prec', 'Precision\n(ALT calls correct)', 100)]
for ax, scope, title in [(b1, 'genome-wide', 'A   Genome-wide (MAF 1–10%)'),
                         (b2, 'gene regions', 'B   Annotated gene regions')]:
    q = t2[(scope, 'Q-Imputer w25')]; g = t2[(scope, 'Beagle 5.4')]
    ys = np.arange(3)[::-1]
    for k, (key, lab, sc) in enumerate(met):
        yq, yg = ys[k] + 0.13, ys[k] - 0.13
        vq, vg = sc * float(q[key]), sc * float(g[key])
        ax.plot([min(vq, vg), max(vq, vg)], [yq, yg], color=GREY, lw=1.2, zorder=2)
        ax.plot(vq, yq, 'o', ms=13, color=RED, zorder=4)
        ax.plot(vg, yg, 'o', ms=13, color=SLATE, zorder=4)
        ax.annotate(f'{vq:.3g}%', (vq, yq), xytext=(0, 12), textcoords='offset points',
                    ha='center', fontsize=9.5, fontweight='bold', color=RED)
        ax.annotate(f'{vg:.3g}%', (vg, yg), xytext=(0, -21), textcoords='offset points',
                    ha='center', fontsize=9.5, fontweight='bold', color=SLATE)
    ax.set_xscale('log'); ax.set_xlim(0.02, 200)
    ax.set_yticks(ys); ax.set_yticklabels([m[1] for m in met], fontsize=10)
    ax.set_ylim(-0.75, 2.85)
    ax.set_xlabel('Rate or precision (%) — logarithmic scale', fontsize=10.5)
    ax.grid(axis='x', alpha=.3, zorder=0)
    ax.set_title(title, fontsize=11.5, fontweight='bold', loc='left')
b1.plot([], [], 'o', color=RED, ms=10, label='QuImputer (w = 25)')
b1.plot([], [], 'o', color=SLATE, ms=10, label='Beagle 5.4')
b1.legend(fontsize=9.5, loc='upper right', framealpha=.95)
plt.tight_layout()
for e in ('png', 'pdf'):
    fig.savefig(os.path.join(OUTD, f'Figure3_rare_alt_profile.{e}'), dpi=300, bbox_inches='tight')
print('Figure 3 saved')


# ================= Supplementary Figure S1 =================
rows = list(csv.DictReader(open(PO + 'figS1_per_chr_maf_binned_w25.csv')))
chrs = [r['Chr'] for r in rows]
bins = [('MAF_lt1', '<1% (very rare)'), ('MAF_1_5', '1-5% (rare)'),
        ('MAF_5_10', '5-10% (low-frequency)'), ('MAF_10_50', '10-50% (common)')]
# 염색체별 all-REF 기준선은 S6 원자료에서
s6 = {r['chrom']: r for r in csv.DictReader(open(PO + 'per_chromosome_w25.tsv'), delimiter='\t')}
BL = {'MAF_lt1': 'allref_MAF_lt1', 'MAF_1_5': 'allref_MAF_1_5',
      'MAF_5_10': 'allref_MAF_5_10', 'MAF_10_50': 'allref_MAF_10_50'}

fig, axes = plt.subplots(2, 2, figsize=(12.8, 7.6))
x = np.arange(12); w = 0.36
for ax, (key, title) in zip(axes.ravel(), bins):
    qi = [float(r[key + '_QI']) for r in rows]
    bg = [float(r[key + '_BG']) for r in rows]
    bl = [float(s6[c][BL[key]]) for c in chrs]
    ax.bar(x - w/2, qi, w, color=RED, label='QuImputer (w = 25)', zorder=3)
    ax.bar(x + w/2, bg, w, color=SLATE, label='Beagle 5.4', zorder=3)
    for i, b in enumerate(bl):
        ax.plot([i - 0.47, i + 0.47], [b, b], color='k', ls='--', lw=1.5, zorder=5,
                label='all-REF baseline' if i == 0 else None)
    lo = min(min(qi), min(bg), min(bl)); hi = max(max(qi), max(bg), max(bl))
    pad = (hi - lo) * 0.35 + 0.002
    ax.set_ylim(lo - pad, hi + pad * 0.6)
    ax.set_xticks(x); ax.set_xticklabels([c.replace('chr', '') for c in chrs], fontsize=9)
    ax.set_xlabel('Chromosome', fontsize=10)
    ax.set_ylabel('Binary concordance', fontsize=10)
    ax.grid(axis='y', alpha=.3, zorder=0)
    ax.set_title(title, fontsize=11, fontweight='bold', loc='left')
axes[0, 0].legend(fontsize=8.5, loc='lower right', framealpha=.95)
plt.tight_layout()
for e in ('png', 'pdf'):
    fig.savefig(os.path.join(OUTD, f'FigureS1_per_chromosome_concordance.{e}'),
                dpi=300, bbox_inches='tight')
print('Figure S1 saved')

# ================= Supplementary Figure S2 =================
g = list(csv.DictReader(open(PO + 'figS2_gene_type2_diff.csv')))
d = np.array([float(r['fnr_diff']) for r in g]) * 100     # QI - Beagle, percentage points
n = len(d)
qi_lower = int((d < -0.1).sum()); bg_lower = int((d > 0.1).sum())
tied = n - qi_lower - bg_lower

fig, (a1, a2) = plt.subplots(1, 2, figsize=(12.6, 4.9),
                             gridspec_kw={'width_ratios': [1.35, 1]})
bw = 0.5
edges = np.arange(-30, 30 + bw, bw)
a1.hist(d, bins=edges, color=GREY, zorder=3)
a1.hist(d[d < -0.1], bins=edges, color=RED, zorder=4, label='QuImputer lower Type II error')
a1.hist(d[d > 0.1], bins=edges, color=SLATE, zorder=4, label='Beagle lower Type II error')
a1.axvline(0, color='k', lw=1.2, zorder=5)
a1.set_yscale('log')
a1.set_xlim(-25, 25)
a1.set_xlabel('Type II error difference, QuImputer - Beagle (percentage points)', fontsize=10.5)
a1.set_ylabel('Genes (log scale)', fontsize=10.5)
a1.grid(alpha=.3, zorder=0)
a1.legend(fontsize=9, loc='upper left', framealpha=.95)
a1.set_title('A   Distribution across %s genes' % f'{n:,}', fontsize=11.5,
             fontweight='bold', loc='left')
a1.text(0.6, a1.get_ylim()[1] * 0.25, 'negative = QuImputer\nmisses fewer carriers',
        fontsize=8.6, color='#5A6B73', style='italic')

cats = ['QuImputer\nlower', 'Tied\n(within 0.1 pp)', 'Beagle\nlower']
vals = [qi_lower, tied, bg_lower]
cols = [RED, GREY, SLATE]
bars = a2.bar(np.arange(3), vals, width=.6, color=cols, zorder=3)
for i, v in enumerate(vals):
    a2.annotate(f'{v:,}\n({100*v/n:.2f}%)', (i, v), xytext=(0, 5),
                textcoords='offset points', ha='center', fontsize=10, fontweight='bold')
a2.set_xticks(np.arange(3)); a2.set_xticklabels(cats, fontsize=10)
a2.set_ylabel('Genes', fontsize=10.5)
a2.set_ylim(0, max(vals) * 1.22)
a2.grid(axis='y', alpha=.3, zorder=0)
a2.set_title('B   Gene-level tally', fontsize=11.5, fontweight='bold', loc='left')
plt.tight_layout()
for e in ('png', 'pdf'):
    fig.savefig(os.path.join(OUTD, f'FigureS2_gene_type2_difference.{e}'),
                dpi=300, bbox_inches='tight')
print('Figure S2 saved | QI %d (%.2f%%) / tied %d (%.2f%%) / BG %d (%.2f%%)'
      % (qi_lower, 100*qi_lower/n, tied, 100*tied/n, bg_lower, 100*bg_lower/n))


import subprocess
subprocess.run([sys.executable, os.path.join(os.path.dirname(__file__), 'make_revised_figures.py'), ROOT], check=True)
print('Figures 2, 3, 4, S1 and S2 regenerated from archive summaries')
```
