## Supplementary data and analysis code for "QuImputer: an LD-informed QUBO/Ising formulation of genotype imputation with quantum-hardware execution": make_figure1_20261007.py

```
"""Draw Figure 1 as a vector PDF. Usage: python make_figure1.py output.pdf

Requires ReportLab and Unicode-capable TrueType fonts. QUIMPUTER_FONT_DIR may
point to Arial Unicode.ttf + Arial Bold.ttf, or DejaVuSans.ttf + DejaVuSans-Bold.ttf.
Standard macOS and Linux font directories are tried automatically.
"""
from pathlib import Path
import os, sys, math
from reportlab.pdfgen import canvas
from reportlab.pdfbase import pdfmetrics
from reportlab.pdfbase.ttfonts import TTFont
from reportlab.lib.colors import HexColor
from reportlab.lib.styles import ParagraphStyle
from reportlab.platypus import Paragraph

def make(out):
    candidates=[Path(os.environ['QUIMPUTER_FONT_DIR'])] if 'QUIMPUTER_FONT_DIR' in os.environ else [Path('/System/Library/Fonts/Supplemental'),Path('/usr/share/fonts/truetype/dejavu')]
    normal=bold=None
    for fontdir in candidates:
        for a,b in [('Arial Unicode.ttf','Arial Bold.ttf'),('DejaVuSans.ttf','DejaVuSans-Bold.ttf')]:
            if (fontdir/a).exists() and (fontdir/b).exists():normal,bold=fontdir/a,fontdir/b;break
        if normal:break
    if normal is None:raise RuntimeError('Set QUIMPUTER_FONT_DIR to a Unicode-capable Arial or DejaVu TrueType font directory.')
    pdfmetrics.registerFont(TTFont('Diagram',str(normal)))
    pdfmetrics.registerFont(TTFont('DiagramBold',str(bold)))
    pdfmetrics.registerFontFamily('Diagram',normal='Diagram',bold='DiagramBold')
    c=canvas.Canvas(str(out),pagesize=(900,510))
    c.setTitle('QuImputer QUBO Ising genotype imputation workflow')
    ink='#203040'; blue='#2679B9'; green='#41824B'; orange='#C57918'; pink='#A94D72'
    def label(x,y,t,w,size=13,color=ink):
        t=t.replace('<sub>','<sub size="8">').replace('<super>','<super size="8">')
        st=ParagraphStyle('p',fontName='Diagram',fontSize=size,leading=size*1.27,textColor=HexColor(color),alignment=1)
        p=Paragraph(t,st);_,h=p.wrap(w,300);p.drawOn(c,x,y-h)
    def box(x,y,w,h,t,stroke,fill,size=13):
        t=t.replace('<sub>','<sub size="8">').replace('<super>','<super size="8">')
        c.setStrokeColor(HexColor(stroke));c.setFillColor(HexColor(fill));c.setLineWidth(1.7)
        c.roundRect(x,y,w,h,6,stroke=1,fill=1)
        st=ParagraphStyle('p',fontName='Diagram',fontSize=size,leading=size*1.27,textColor=HexColor(ink),alignment=1)
        p=Paragraph(t,st);_,ph=p.wrap(w-20,h-10);assert ph<h-7,(t,ph,h)
        p.drawOn(c,x+10,y+(h-ph)/2)
    def line(points,color=ink,arrow=True):
        c.setStrokeColor(HexColor(color));c.setFillColor(HexColor(color));c.setLineWidth(1.5)
        p=c.beginPath();p.moveTo(*points[0])
        for x,y in points[1:]:p.lineTo(x,y)
        c.drawPath(p)
        if arrow:
            x,y=points[-1];a=math.atan2(y-points[-2][1],x-points[-2][0]);s=6
            p=c.beginPath();p.moveTo(x,y);p.lineTo(x-s*math.cos(a-.5),y-s*math.sin(a-.5));p.lineTo(x-s*math.cos(a+.5),y-s*math.sin(a+.5));p.close();c.drawPath(p,fill=1,stroke=0)
    label(30,504,'<b>QuImputer QUBO/Ising genotype-imputation workflow</b>',840,18)
    box(30,405,210,66,'<b>Masked VCF genotypes</b><br/>Observed and missing states',blue,'#EEF5FB')
    box(280,405,250,66,'<b>Population statistics</b><br/>Allele frequencies and signed LD',green,'#F0F6EF')
    box(650,405,220,66,'<b>Beagle 5.4</b><br/>HMM phasing and imputation',ink,'#EFF2F5')
    box(30,290,210,76,'<b>Observed markers</b><br/>Fixed states contribute<br/>to local fields',blue,'#EEF5FB')
    box(280,276,250,96,'<b>Reduced QUBO/Ising objective</b><br/>Missing SNPs are variables<br/>H = Σ h<sub>i</sub>s<sub>i</sub> + Σ<sub>i&lt;j</sub> J<sub>ij</sub>s<sub>i</sub>s<sub>j</sub><br/>Joint optimization within each window',orange,'#FFF6E8',12.5)
    line([(240,438),(280,438)],blue)
    line([(135,405),(135,366)],blue)
    line([(240,328),(280,328)],blue)
    line([(405,405),(405,372)],green)
    # Both pipelines receive the same masked matrix in the primary benchmark.
    line([(135,471),(135,481),(760,481),(760,471)],ink)
    label(547,401,'Same masked input',92,10)
    label(32,266,'Same objective, alternative solvers',235,11)
    line([(405,276),(405,244)],orange,False)
    line([(150,244),(750,244)],orange,False)
    for x in [150,450,750]:line([(x,244),(x,225)],orange)
    box(30,153,240,72,'<b>Exact classical search</b><br/>NumPy exhaustive enumeration<br/>min<sub>x</sub> x<super>T</super>Qx',orange,'#FFF6E8')
    box(330,153,240,72,'<b>QAOA simulation</b><br/>Cost layer → mixer layer<br/>Circuit sampling',pink,'#FBEFF4')
    box(630,153,240,72,'<b>Selected IBM QPU executions</b><br/>QAOA circuits on IBM Fez<br/>156-qubit Heron processor',pink,'#FBEFF4')
    for x in [150,450,750]:line([(x,153),(x,137)],orange,False)
    line([(150,137),(750,137)],orange,False)
    line([(405,137),(405,122)],orange)
    box(280,67,250,55,'<b>QuImputer assignments</b><br/>REF or ALT-carrier state',pink,'#FBEFF4')
    box(650,57,220,77,'<b>Evaluation against known truth</b><br/>Binary concordance, ALT-carrier<br/>metrics and hard-call dosage R²',green,'#F0F6EF',12)
    line([(530,94),(650,94)],pink)
    line([(870,438),(888,438),(888,111),(870,111)],ink)
    label(30,40,'QAOA state for one layer: |ψ〉 = exp(−iβH<sub>M</sub>) exp(−iγH<sub>C</sub>) |+〉<super>⊗n</super>',840,13)
    c.showPage();c.save()

if __name__=='__main__':make(sys.argv[1])
```
