## Supplementary data and analysis code for "QuImputer: an LD-informed QUBO/Ising formulation of genotype imputation with quantum-hardware execution": make_figure6.py

```
"""Render Figure 6 from the published September 2026 execution summary.

Usage: python scripts/make_figure6.py /path/to/QuImputer_supplementary_data
This is a plotting operation only; it performs no experiment or inference.
"""
from pathlib import Path
import csv
import sys
import matplotlib
matplotlib.use('Agg')
import matplotlib.pyplot as plt
import numpy as np

root = Path(sys.argv[1] if len(sys.argv) > 1 else '.')
with (root/'quantum_hardware/selected_validation_timings.csv').open() as f:
    records = {(int(r['problem']),r['backend']):r for r in csv.DictReader(f)}
plt.rcParams.update({'font.family':'DejaVu Sans','font.size':9,'axes.spines.top':False,
                     'axes.spines.right':False,'pdf.fonttype':42,'ps.fonttype':42})
fig,axes=plt.subplots(1,2,figsize=(7.0,3.0))
x=np.arange(5); width=.34
for ax,field,label,panel in [(axes[0],'exact_rate_percent','Pooled exact-optimum shots (%)','A'),
                              (axes[1],'elapsed_wall_s','Elapsed wall time (s)','B')]:
    for bi,(backend,color,name) in enumerate([('aer_simulator','#4477AA','Aer'),('ibm_fez','#CC6677','IBM Fez')]):
        vals=[float(records[(i,backend)][field]) for i in range(1,6)]
        ax.bar(x+(bi-.5)*width,vals,width,label=name,color=color)
    ax.set_xticks(x,[f'{i}\n({records[(i,"aer_simulator")]["n_qubits"]} q)' for i in range(1,6)])
    ax.set_xlabel('Problem (active qubits)')
    ax.set_ylabel(label)
    ax.text(0,1.035,panel,transform=ax.transAxes,fontweight='bold',fontsize=11)
    ax.yaxis.grid(True,color='#E4E4E4',linewidth=.6)
    ax.set_axisbelow(True)
axes[0].set_ylim(0,3.1); axes[1].set_ylim(0,170)
handles,labels=axes[0].get_legend_handles_labels()
fig.legend(handles,labels,frameon=False,fontsize=8,loc='upper center',ncol=2,bbox_to_anchor=(.5,1.0))
fig.subplots_adjust(left=.08,right=.98,bottom=.25,top=.83,wspace=.42)
for ext in ['png','pdf']:
    fig.savefig(root/f'figures/Figure6_validation_rerun.{ext}',dpi=300)
plt.close(fig)
```
