## Supplementary data and analysis code for "QuImputer: an LD-informed QUBO/Ising formulation of genotype imputation with quantum-hardware execution": make_figure7_20261006.py

```
"""Generate publication Figure 7 from the supplied metrics JSON (ReportLab).
Usage: python make_figure7.py paper_metrics.json Figure7_hardware_20261006.pdf
"""
import json,math,sys
from pathlib import Path
from reportlab.pdfgen import canvas
from reportlab.lib.colors import HexColor,Color,black,white

def make(src,out):
    rows=json.loads(Path(src).read_text())['problems'];W,H=504,410;c=canvas.Canvas(str(out),pagesize=(W,H))
    blue=HexColor('#2679b9');orange=HexColor('#db632d');gray=HexColor('#666666');grid=HexColor('#dddddd')
    left,right=58,490;xs=[left+17+i*35.0 for i in range(12)]
    def text(x,y,s,size=8,color=black,align='left',bold=False):
        c.setFillColor(color);c.setFont('Helvetica-Bold' if bold else 'Helvetica',size)
        (c.drawCentredString if align=='center' else c.drawRightString if align=='right' else c.drawString)(x,y,s)
    def line(x1,y1,x2,y2,color=gray,width=.6,dash=None):
        c.setStrokeColor(color);c.setLineWidth(width);c.setDash(dash or []);c.line(x1,y1,x2,y2);c.setDash([])
    def marker(x,y,color,kind,fill=True):
        c.setStrokeColor(color);c.setFillColor(color if fill else white);c.setLineWidth(1.1)
        if kind=='circle':c.circle(x,y,3.2,stroke=1,fill=1)
        else:
            p=c.beginPath();p.moveTo(x,y+4);p.lineTo(x+4,y);p.lineTo(x,y-4);p.lineTo(x-4,y);p.close();c.drawPath(p,stroke=1,fill=1)
    def labelvertical(x,y,s):
        c.saveState();c.translate(x,y);c.rotate(90);text(0,0,s,8,align='center');c.restoreState()
    ay0,ay1=244,369
    def ya(v):return ay0+(math.log10(v)+3)/(math.log10(2500)+3)*(ay1-ay0)
    text(left,392,'A  Exact-optimum shots across the parameter grid',10,bold=True)
    for v in [.001,.01,.1,1,10,100,1000]:
        y=ya(v);line(left,y,right,y,grid);text(left-7,y-2.6,f'{v:g}',7.5,align='right')
    line(left,ay0,left,ay1);line(left,ay0,right,ay0)
    c.setFillColor(HexColor('#f3f3f3'));c.rect(left,218,right-left,14,fill=1,stroke=0)
    text(left-7,222,'none',7.5,align='right')
    labelvertical(14,300,'Number of exact-optimum shots (log scale)')
    for x,r in zip(xs,rows):
        marker(x-4,ya(r['ideal_expected_optimum_shots']),blue,'circle')
        if r['optimum_shots']:marker(x+4,ya(r['optimum_shots']),orange,'diamond')
        else:marker(x+4,225,orange,'diamond',False)
    marker(263,383,blue,'circle');text(271,380,'Noiseless expectation',7.5)
    marker(379,383,orange,'diamond');text(387,380,'IBM Fez',7.5)
    text(left,202,'Open diamonds: zero observed. Noiseless expectations are exact calculations.',7.2,color=gray)
    by0,by1=58,169
    def yb(v):return by0+(100*v-70)/31*(by1-by0)
    text(left,183,'B  Selected-output concordance with known biological truth',10,bold=True)
    for v in [.7,.8,.9,1]:
        y=yb(v);line(left,y,right,y,grid);text(left-7,y-2.6,f'{100*v:g}',7.5,align='right')
    line(left,by0,left,by1);line(left,by0,right,by0);labelvertical(14,111,'Matching known-truth variables (%)')
    for x,r in zip(xs,rows):
        a=r['all_ref_truth_matches']/r['truth_denominator'];b=r['truth_matches']/r['truth_denominator']
        line(x-7,yb(a),x+7,yb(a),blue,1.8)
        marker(x,yb(b),orange,'diamond')
        text(x,44,r['problem_id'],7,align='center')
    split=(xs[-1]+xs[-2])/2
    for bottom,top in [(218,369),(by0,by1)]:line(split,bottom,split,top,grid,1,dash=[2,2])
    line(236,16,248,16,blue,1.8);text(253,13,'all-REF / exact / local-field-only',7)
    marker(415,16,orange,'diamond');text(423,13,'selected QPU',7)
    text(left,29,'A: 25 circuits, 102,400 shots per instance     B120: 100 circuits, 409,600 shots',7.1,color=gray)
    c.showPage();c.save()
if __name__=='__main__':make(sys.argv[1],sys.argv[2])
```
