## Supplementary data and analysis code for "QuImputer: an LD-informed QUBO/Ising formulation of genotype imputation with quantum-hardware execution": make_revised_figures.py

```
"""Regenerate Figures 2 and 4 from archive source summaries.

Dependencies: reportlab and Poppler pdftoppm. Run with the archive root as argument.
PDF figures remain vector graphics; PNGs are rasterized at 250 dpi for Word.
"""
import csv,json,math,sys,subprocess,shutil,os
from pathlib import Path
from reportlab.pdfgen import canvas
from reportlab.lib.colors import HexColor,white,black

ROOT=Path(sys.argv[1]) if len(sys.argv)>1 else Path(__file__).resolve().parent.parent
OUT=ROOT/'figures';OUT.mkdir(exist_ok=True)
RED='#B64035';BLUE='#3D5A6C';GRAY='#9AA5AB';GRID='#DDE1E5';INK='#17212B'
WIDTH,HEIGHT=760,315

def txt(c,x,y,s,size=10,color=INK,bold=False,align='left'):
    c.setFillColor(HexColor(color));c.setFont('Helvetica-Bold' if bold else 'Helvetica',size)
    getattr(c,{'left':'drawString','center':'drawCentredString','right':'drawRightString'}[align])(x,y,str(s))
def line(c,x1,y1,x2,y2,color=GRAY,width=.6,dash=None):
    c.setStrokeColor(HexColor(color));c.setLineWidth(width);c.setDash(dash or [])
    c.line(x1,y1,x2,y2);c.setDash([])
def bar(c,x,y,w,h,color):
    c.setFillColor(HexColor(color));c.rect(x,y,w,max(h,.35),stroke=0,fill=1)
def dot(c,x,y,color=RED,triangle=False,r=3.1):
    c.setFillColor(HexColor(color));c.setStrokeColor(HexColor(color))
    if triangle:
        p=c.beginPath();p.moveTo(x-r,y+r);p.lineTo(x+r,y+r);p.lineTo(x,y-r);p.close();c.drawPath(p,stroke=0,fill=1)
    else:c.circle(x,y,r,stroke=0,fill=1)
def poly(c,pts,color=RED,width=1.3,dash=None):
    for (x,y),(a,b) in zip(pts,pts[1:]):line(c,x,y,a,b,color,width,dash)
def canvas_for(name):
    c=canvas.Canvas(str(OUT/(name+'.pdf')),pagesize=(WIDTH,HEIGHT));c.setTitle(name);c.setAuthor('QuImputer authors');return c
def done(c,name):
    c.showPage();c.save()
    pop=os.environ.get('QUIMPUTER_PDFTOPPM') or shutil.which('pdftoppm')
    if not pop: raise RuntimeError('pdftoppm is required for PNG output')
    subprocess.run([pop,'-png','-singlefile','-r','250',str(OUT/(name+'.pdf')),str(OUT/name)],check=True,capture_output=True)
    print(name)
def axes(c,x,y,w,h,ticks,lo,hi,label,transform=lambda v:v):
    f=lambda v:y+h*(transform(v)-transform(lo))/(transform(hi)-transform(lo))
    for v,lab in ticks:
        yy=f(v);line(c,x,yy,x+w,yy,GRID);txt(c,x-8,yy-3,lab,8.5,align='right')
    line(c,x,y,x,y+h,INK);line(c,x,y,x+w,y,INK)
    c.saveState();c.translate(x-38,y+h/2);c.rotate(90);txt(c,0,0,label,9.5,align='center');c.restoreState()
    return f

def fig2():
    name='Figure2_maf_concordance';c=canvas_for(name)
    rows=list(csv.DictReader(open(ROOT/'primary_observed/G2_table1.csv')))[:4]
    labs=['<1%','1-5%','5-10%','10-50%']
    txt(c,48,288,'A   Concordance and all-REF baseline',12,bold=True)
    txt(c,422,288,'B   Difference from all-REF',12,bold=True)
    f=axes(c,52,82,286,172,[(.7,'.70'),(.8,'.80'),(.9,'.90'),(1,'1.00')],.65,1.015,'Binary concordance')
    for i,r in enumerate(rows):
        x=83+i*69
        for off,key,col in [(-20,'qi_w25',RED),(2,'beagle',BLUE)]:
            v=float(r[key]);bar(c,x+off,f(.65),18,f(v)-f(.65),col)
        line(c,x-24,f(float(r['all_ref_baseline'])),x+25,f(float(r['all_ref_baseline'])),INK,1,[3,2])
        txt(c,x,65,labs[i],9,align='center')
    sym=lambda v: math.copysign(abs(v)/.05 if abs(v)<=.05 else 1+math.log(abs(v)/.05),v)
    g=axes(c,434,82,286,172,[(-.1,'-0.1'),(0,'0'),(.1,'0.1'),(1,'1'),(10,'10')],-.4,25,'Excess concordance (pp, symmetric log)',sym)
    line(c,434,g(0),720,g(0),INK,.9)
    for i,r in enumerate(rows):
        x=465+i*69
        for off,key,col in [(-20,'qi_w25_excess_pp',RED),(2,'beagle_excess_pp',BLUE)]:
            v=float(r[key]);yy=g(v);bar(c,x+off,min(yy,g(0)),18,abs(yy-g(0)),col)
            label=f'{v:+.4f}' if abs(v)<1 else f'{v:+.2f}'
            txt(c,x+off+9,yy+(6 if v>=0 else -12)-(10 if i==0 and off==2 else 0),label,7.5,col,align='center')
        txt(c,x,65,labs[i],9,align='center')
    txt(c,195,48,'Minor allele frequency',9,align='center');txt(c,577,48,'Minor allele frequency',9,align='center')
    bar(c,64,23,10,7,RED);txt(c,80,22,'QuImputer w = 25',9)
    bar(c,211,23,10,7,BLUE);txt(c,227,22,'Beagle 5.4',9)
    line(c,325,27,345,27,INK,1,[3,2]);txt(c,350,22,'All-REF baseline',9)
    txt(c,720,22,'Primary observed-data evaluation',8.5,align='right')
    done(c,name)

def fig4():
    name='Figure4_window_constraint';c=canvas_for(name)
    d=json.loads((ROOT/'mechanism/W1_alt_neighbour_raw.json').read_text())
    bins=['MAF<1%','MAF1-5%','MAF5-10%','MAF10-50%'];labs=['<1%','1-5%','5-10%','10-50%']
    txt(c,48,288,'A   Local LD support for masked ALT carriers',11.5,bold=True)
    txt(c,48,272,'First 100,000 SNPs of chr01; 25-SNP blocks',9)
    f=axes(c,52,82,260,168,[(0,'0'),(.5,'0.5'),(1,'1.0'),(1.5,'1.5')],0,1.8,'Mean LD partners with r² > 0.01')
    pts=[]
    for i,b in enumerate(bins):
        v=d['ldpart'][b];n=sum(v);m=sum(j*k for j,k in enumerate(v))/n
        x=85+i*63;bar(c,x-15,82,30,f(m)-82,RED);txt(c,x,f(m)+7,f'{m:.3f}',10,RED,True,'center')
        a=d['hist_alt'][b];z=100*a[0]/sum(a);yy=82+(z-90)/12*168;pts.append((x,yy))
        dot(c,x,yy,BLUE);txt(c,x,yy-16,f'{z:.1f}%',8.5,BLUE,align='center')
        txt(c,x,65,labs[i],9,align='center')
    poly(c,pts,BLUE,1.2,[3,2])
    for v in [90,95,100]:txt(c,318,82+(v-90)/12*168-3,f'{v}%',8,BLUE)
    txt(c,182,47,'Minor allele frequency',9,align='center')
    txt(c,422,288,'B   Distance to the nearest observed ALT',11.5,bold=True)
    txt(c,422,272,'Whole chr01; no LD requirement for the neighbour',9)
    g=axes(c,432,82,288,168,[(0,'0'),(25,'25'),(50,'50'),(75,'75'),(100,'100')],0,100,'Evaluated carriers within distance (%)')
    xx=lambda k:432+288*(math.log10(k)-math.log10(10))/(math.log10(6000)-math.log10(10))
    pts=[]
    for k,v in sorted((int(k),v) for k,v in d['nearest_within'].items()):
        pts.append((xx(k),g(100*v/d['nearest_n'])))
    poly(c,pts,RED,1.5)
    for x,y in pts:dot(c,x,y,RED)
    for k in [12,50,200,1000,5000]:txt(c,xx(k),65,str(k),8.5,align='center')
    for k,dy in [(12,8),(100,9),(500,-15)]:
        v=100*d['nearest_within'][str(k)]/d['nearest_n'];txt(c,xx(k)+4,g(v)+dy,f'{v:.2f}%',8.5,RED)
    txt(c,576,47,'Distance in SNPs (log scale)',9,align='center')
    line(c,52,25,69,25,BLUE,1.2,[3,2]);txt(c,76,22,'No observed ALT-carrier LD partner (right scale)',8.5,BLUE)
    txt(c,720,22,'n = 2,934,770; median = 74 SNPs',9,align='right')
    done(c,name)

if __name__=='__main__':fig2();fig4()
```
